# Myosin-X and talin modulate integrin activity at filopodia tips

**DOI:** 10.1101/2020.05.05.078733

**Authors:** Mitro Miihkinen, Max L.B. Grönloh, Helena Vihinen, Eija Jokitalo, Benjamin T. Goult, Johanna Ivaska, Guillaume Jacquemet

**Author notes:** **Correspondence to**: Guillaume Jacquemet or Johanna Ivaska. Current address: Molecular Cell Biology lab, Department of Molecular Cellular Hemostasis, Sanquin Research and Landsteiner Laboratory, 1066 CX Amsterdam, The Netherlands.

## Abstract

Filopodia assemble unique integrin-adhesion complexes to sense the extracellular matrix. However, the mechanisms of integrin localization and regulation in filopodia are poorly defined. Here, we observed that active integrins accumulate at the tip of myosin-X(MYO10)-positive filopodia while inactive integrins are uniformly distributed. RNAi depletion of 10 integrin activity modulators identified talin as the principal integrin activator in filopodia. Deletion of the MYO10-FERM domain, or mutation of the β1-integrin-binding residues within, revealed MYO10 as facilitating integrin activation but not transport in filopodia. However, MYO10-FERM alone could not activate integrins, potentially due to dual binding to both a- and β-integrin tails. As swapping MYO10-FERM with talin-FERM enabled integrin activation in filopodia, our data indicate that an integrin-binding FERM domain coupled to a myosin motor is a core requirement for integrin activation in filopodia. Therefore, we propose a two-step integrin activation model in filopodia: receptor tethering by MYO10 followed by talin-mediated integrin activation.

## Introduction

Filopodia are actin-rich “antenna-like” protrusions that are responsible for constantly probing the cellular environment composed of neighboring cells and the extracellular matrix (ECM). As such, filopodia contain cell-surface receptors such as integrins, cadherins, and growth factor receptors that can interact with, and interpret, a wide variety of extracellular cues (Jacquemet et al., 2015). Filopodia are especially abundant in cells as they migrate in 3D and in vivo where they contribute to efficient directional migration by probing and remodeling the surrounding ECM (Jacquemet et al., 2013, 2017; Paul et al., 2015).

Filopodia have a unique cytoskeleton composed of tightly packed parallel actin filaments with barbed ends oriented towards the filopodium tip (Mattila and Lappalainen, 2008). This organization allows molecular motors, such as unconventional myosin-X (MYO10), to move towards and accumulate at the tips. By doing so, these molecular motors are thought to transport various proteins, including integrins, along actin filaments to the tips of filopodia (Jacquemet et al., 2015; Arjonen et al., 2014; Berg and Cheney, 2002; Hirano et al., 2011; Zhang et al., 2004). In particular, MYO10 is known to bind directly to the NPxY motif of the β-integrin cytoplasmic tail via its FERM (protein 4.1R, ezrin, radixin, moesin) domain (Zhang et al., 2004). At filopodia tips, integrins assemble a specific adhesion complex that tethers filopodia to the ECM (Alieva et al., 2019; Jacquemet et al., 2019; Gallop, 2019). Filopodia adhesions contain several adhesion proteins including talin, kindlin, and p130Cas but are devoid of the nascent adhesion markers focal adhesion kinase (FAK) and paxillin (Jacquemet et al., 2019), indicating that filopodia adhesions are distinct in their molecular composition from other adhesion types. The subsequent maturation of these filopodia adhesions into nascent and focal adhesions can promote directional cell migration (Hu et al., 2014; Jacquemet et al., 2016, 2019).

Integrin functions are tightly regulated by a conformational switch that modulates ECM binding, often referred to as activation. Integrin extracellular domain conformations can range from a bent to an extended open conformation, where the integrin’s ligand affinity increases with stepwise opening (Conway and Jacquemet, 2019; Sun et al., 2019; Askari et al., 2009). For β1 integrin, this unfolding can be viewed using activation-specific antibodies (Byron et al., 2009). Mechanistically, integrin activity can be finely tuned, from within the cell, by multiple proteins that bind to the integrin cytoplasmic tails (Conway and Jacquemet, 2019; Sun et al., 2019; Askari et al., 2009; Bouvard et al., 2013). For instance, talin, a key integrin activator, can bind to the NPxY motif of the β integrin cytoplasmic tail leading to the physical separation of the integrin a and β cytoplasmic tails and integrin activation. Kindlin, another critical regulator of integrin activity, also binds to β integrin cytoplasmic tails where it cooperates with talin to induce integrin activation (Sun et al., 2019). While it is clear that integrins and integrin signaling are key regulators of filopodia function (Lagarrigue et al., 2015; Jacquemet et al., 2016, 2019; Gallop, 2019), how integrin activity is regulated within filopodia is not fully understood.

Here, we observed that active (high affinity) integrin accumulates at filopodia tips while inactive (unoccupied) integrin localizes throughout the filopodia shaft. We find that integrin activation in filopodia is uncoupled from focal adhesions or the actomyosin machinery but is instead locally regulated by talin and MYO10. Contrary to previous assumptions, the FERM domain of MYO10 is not required to transport integrins to filopodia but instead functions to activate integrins at filopodia tips. As MYO10 contributes to integrin activation at the filopodia tips, but MYO10-FERM alone does not directly activate integrins, our data supports a two-step integrin activation model in filopodia. In this model, MYO10 enables integrin receptor tethering at filopodia tips, which is then followed by talin-mediated integrin activation.

## Results

### Integrin activation occurs at filopodia tips independently of cellular forces and focal adhesions

We and others have previously described the formation of integrin-mediated ECM-sensing adhesions at filopodia tips (Shibue et al., 2012; Jacquemet et al., 2019; Lagarrigue et al., 2015; Alieva et al., 2019; Gallop, 2019). To gain further insights into how integrin activity is regulated in MYO10 filopodia, we first assessed the spatial distribution of high affinity and unoccupied β1 integrin (termed active and inactive integrin, respectively, for simplicity) in U2-OS cells transiently expressing fluorescently-tagged MYO10 using structured illumination microscopy (SIM) (Fig. 1A-C) and scanning electron microscopy (SEM) (Fig. 1D). The average distribution of the β1 integrin species along filopodia was mapped from the SIM and the SEM images revealing enrichment and clustering of active β1 integrins at filopodia tips (Fig. 1B-E). In contrast, inactive β1 integrins were more uniformly distributed along the entire length of the filopodium (Fig. 1B-E). Importantly, this pattern of integrin localization was also recapitulated in endogenous filopodia, forming in actively spreading cells (Fig. 1F and 1G).

**Figure 1.**
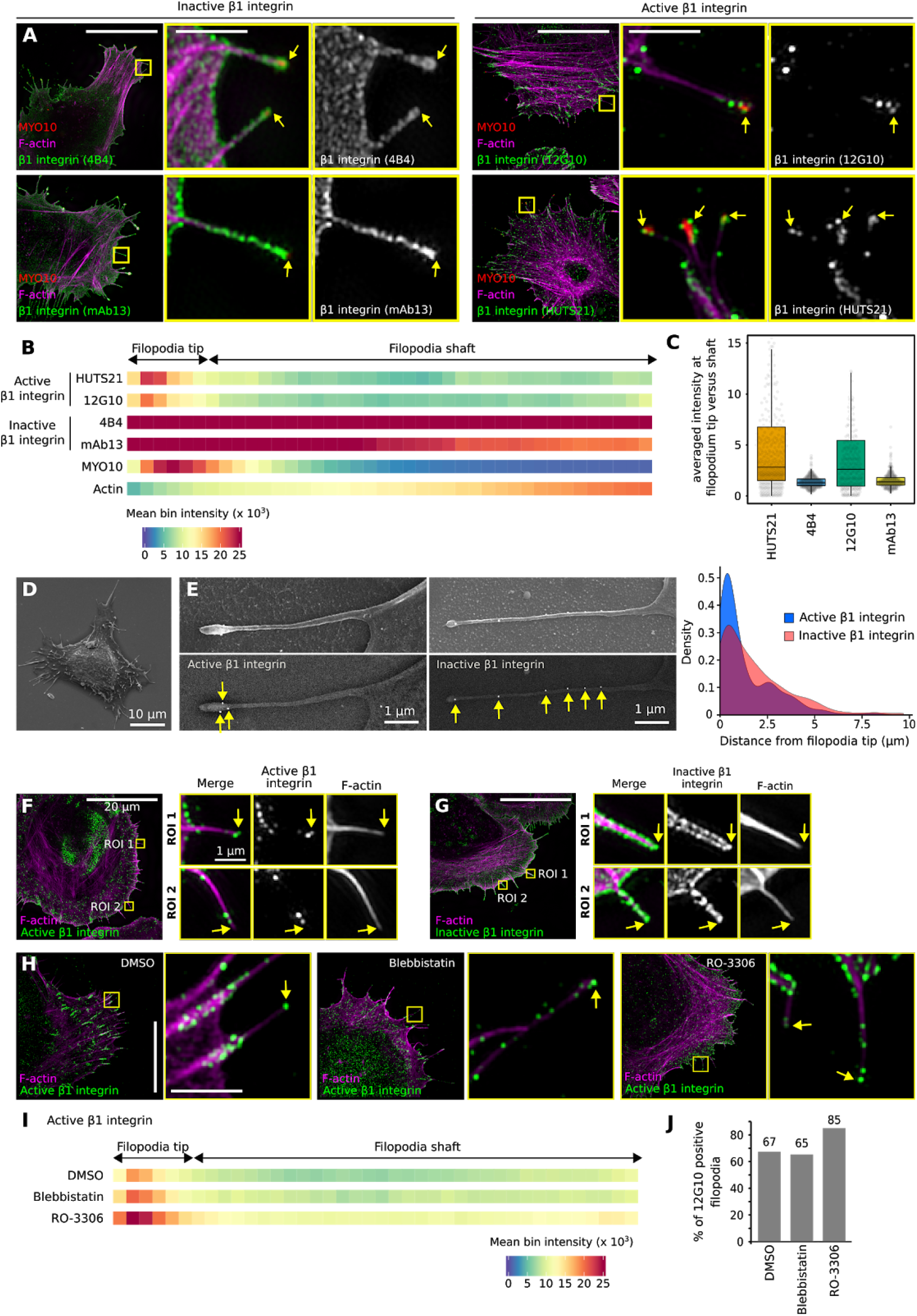
Active integrins accumulate at filopodia tips independently of the cellular forces generated at focal adhesion. **(A-C**) U2-OS cells expressing mScarlet-MYO10 or EGFP-MYO10 were plated on fibronectin for 2 h, stained for active (high-affinity β1 integrin; antibodies 12G10 and HUTS21) or inactive (unoccupied β1 integrin, antibodies 4B4 and mAb13) β1 integrin and F-actin, and imaged using structured illumination microscopy (SIM). Representative maximum intensity projections (MIPs) are displayed. The yellow squares highlight regions of interest (ROIs), which are magnified; yellow arrows highlight filopodia tips; scale bars: (main) 20 μm; (inset) 2 μm. **(B**) Heatmap highlighting the sub-filopodial localization of the proteins stained in A based on their intensity profiles. The segments corresponding to the filopodia tip and shaft are indicated. **(C**) The preferential recruitment of active and inactive β1 integrin to filopodia tips or shafts was assessed by calculating an enrichment ratio (averaged intensity at filopodium tip versus shaft). Results are displayed as Tukey box plots (**B-C)**, See methods for details, MYO10, n = 623 filopodia; F-actin, n = 623 filopodia; HUTS21, n = 538 filopodia; 12G10, n = 329 filopodia; 4B4, n = 413 filopodia; mAb13, n = 369 filopodia; thee biological repeats). (**D-E**) U2-OS cells expressing EGFP-MYO10 were plated on fibronectin for 2 h, stained for active (antibody 12G10) or inactive (antibody 4B4) β1 integrin, and imaged using a scanning electron microscope (SEM). (**D**) A representative image of a whole cell is displayed. (**E**) Representative images of single filopodia are displayed. The upper row was acquired using a secondary electron detector (SED) and the lower row using a backscattered electron detector (vCD). The distance of the two β1 integrin pools (defined by gold particles, highlighted by yellow arrows) from the filopodia tip was measured, and the results displayed as a density plot (4B4 staining, n = 175 gold particles; 12G10 staining, n = 178 gold particles). A bootstrap version of the univariate Kolmogorov-Smirnov test was used for statistical testing. p-value < 0.001. **(F-G**) U2-OS cells were plated on fibronectin for 20 min, stained for active (**E**, antibody 12G10) or inactive (**F**, antibody 4B4) β1 integrin and F-actin, and imaged using SIM. Representative MIPs are displayed. The yellow squares highlight ROIs, which are magnified; yellow arrows highlight filopodia tips; scale bars: (main) 20 μm; (inset) 2 μm. **(H-J**) U2-OS cells expressing EGFP-MYO10 were plated on fibronectin for 1 h and treated for 1 h with 10 μM blebbistatin (myosin II inhibitor), 10 μM RO-3306 (CDK1 inhibitor), or DMSO. Cells were stained for active β1 integrin (antibody 12G10) and F-actin and imaged using SIM. (**H**) Representative MIPs are displayed. The yellow squares highlight ROIs, which are magnified; yellow arrows highlight filopodia tips; scale bars: (main) 20 μm; (inset) 2 μm. (**I**) Heatmap displaying the sub-filopodial localization of active β1 integrin in cells treated with DMSO, blebbistatin or RO-3306 (DMSO, n = 734 filopodia; RO-3306, n = 824 filopodia; blebbistatin, n = 483 filopodia; three biological repeats). (**J**) Bar chart highlighting the percentage of filopodia with detectable levels of active β1 integrin in cells treated with DMSO, blebbistatin or RO-3306 (DMSO, n = 734 filopodia; RO-3306, n = 824 filopodia; blebbistatin, n = 483 filopodia; three biological repeats).

Previous work reported that forces generated by the actomyosin machinery are required for integrin-mediated adhesion at filopodia tips (Alieva et al., 2019). In addition, we observed that filopodia often align to the force generated by focal adhesions (Stubb et al., 2020). Therefore, we investigated whether cellular forces generated by the cell body and transmitted at focal adhesions were responsible for integrin activation at filopodia tips. U2-OS cells expressing fluorescently-tagged MYO10 and adhering to fibronectin were treated with DMSO, a myosin II inhibitor (10 μM blebbistatin) or an established focal adhesion inhibitor (CDK1 inhibitor, 10 μM RO-3306 (Robertson et al., 2015; Jones et al., 2018)). As expected, inhibition of myosin II or CDK1 led to rapid disassembly of focal adhesions (Fig. 1H and S1A). Blebbistatin treatment promoted longer and more numerous filopodia, in line with our earlier report (Stubb et al., 2020), while treatment with the CDK1 inhibitor increased filopodia numbers but not filopodia length (Fig. S1B and S1C). However, no decrease in filopodial integrin activation could be observed when myosin II or CDK1 were inhibited (Fig. 1H and 1I). In contrast, CDK1 inhibition led to an increase in the percentage of filopodia with active integrin at their tips (Fig. 1J). Altogether these data indicate that integrin activation at filopodia tips is regulated independently of cellular forces and focal adhesions. Nevertheless, cellular forces are likely required to induce filopodia adhesion maturation into focal adhesions and for efficient ECM sensing (Alieva et al., 2019; Jacquemet et al., 2019).

### Talin is required to activate β1 integrin at filopodia tips

The enrichment of active β1 integrin at filopodia tips (Fig. 1) indicates that β1 integrin activation is likely to be spatially regulated by one or multiple components of the filopodium-tip complex. We and others have previously reported that several proteins implicated in the regulation of integrin activity, including the integrin activators talins and kindlins as well as the integrin inactivator ICAP-1 (ITGB1BP1), accumulate at filopodia tips where their function remains largely unknown (Lagarrigue et al., 2015; Jacquemet et al., 2016). In addition, we previously reported that enhanced integrin activity often correlates with increased filopodia numbers and stability (Jacquemet et al., 2016). Therefore, we set-up a microscopy-based siRNA screen to test the contribution of 10 known integrin activity regulators on filopodia formation. Each target was silenced with two independent siRNA oligos in U2-OS cells stably expressing MYO10-GFP (Fig. 2A). The effect on MYO10-positive filopodia was scored and the silencing efficiency of each siRNA was validated by qPCR (Fig. S1D) or western blot (Fig. S1E-F). Of the 10 integrin regulators, only talin (combined TLN1 and TLN2) silencing significantly reduced filopodia numbers. As kindlin-2 (FERMT2) is a major regulator of integrin activity (Theodosiou et al., 2016) and FERMT2 localizes to filopodia tips (Jacquemet et al., 2019), we were surprised that FERMT2 silencing did not impact filopodia. To validate this further, we imaged filopodia dynamics in cells silenced for both FERMT1 and FERMT2 (over 90% silencing efficiency). There was no effect on filopodia number or dynamics suggesting that kindlins are not directly required to support filopodia formation or adhesion under the conditions tested (Fig. S1F-H).

**Figure 2.**
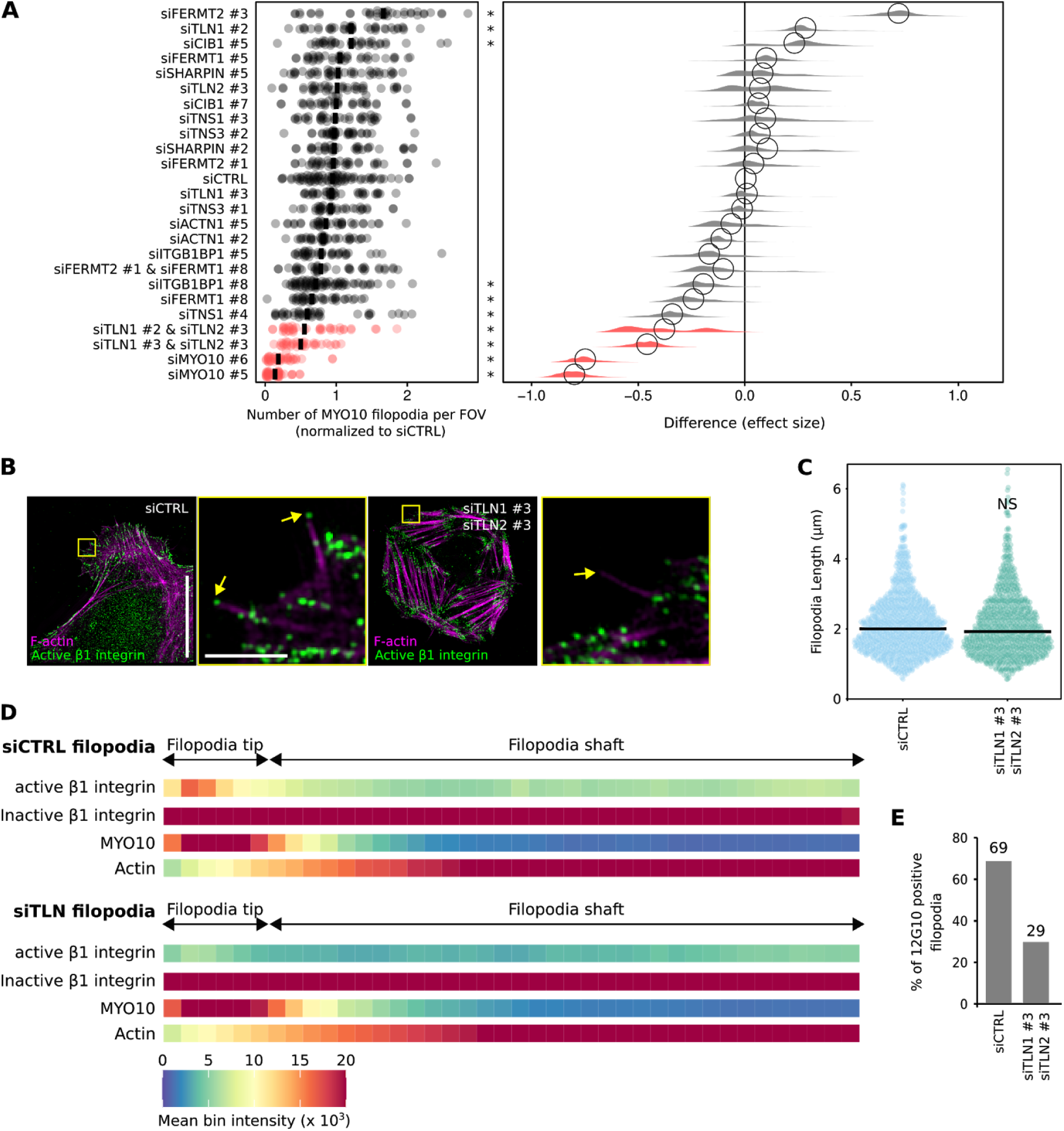
Talin regulates integrin activity at filopodia tips. **(A**) RNAi screen of known integrin activity regulators. The indicated genes were silenced individually or together in U2-OS cells stably expressing EGFP-MYO10 using two independent siRNA oligos per gene. Cells were seeded on fibronectin-coated glass-bottom 96-well plates for 2 h and samples were fixed and stained. Samples were imaged using a spinning-disk confocal microscope and the number of filopodia per field of view (FOV) was quantified automatically using ImageJ (see Methods for details). Results are displayed as dot plots. In addition, the effect size was calculated using bootstrapping to resample the median values for each of the conditions using PlotsOfDifferences (Goedhart, 2019). * p-value < 0.05. **(B-E)** Partially TLN1- and TLN2-silenced U2-OS cells transiently expressing EGFP-MYO10 were plated on fibronectin, stained for active (antibody 12G10) or inactive (antibody mAb13) β1 integrin and F-actin, and imaged using SIM. (**B**) Representative MIPs are displayed. The yellow squares highlight regions of interest (ROIs), which are magnified; yellow arrows highlight filopodia tips; scale bars: (main) 20 μm; (inset) 2 μm. **(C)** Quantification of filopodia length, from the SIM images are displayed as dot plots where the median is highlighted (siCTRL, n = 1086 filopodia; siTLN1 #3 and siTLN2 #3, n = 890 filopodia; three biological repeats). **(D**) Heatmap highlighting the sub-filopodial localisation of the proteins as indicated based on their intensity profiles (siCTRL filopodia: MYO10, n = 799 filopodia; F-actin, n = 799 filopodia; active β1 integrin, n = 799 filopodia; inactive β1 integrin, n = 878 filopodia. siTLN filopodia: MYO10, n = 889 filopodia; F-actin, n = 889 filopodia; active β1 integrin, n = 889 filopodia; inactive β1 integrin, n = 802 filopodia; three biological repeats). (**E**) Bar chart highlighting the percentage of filopodia with detectable levels of active β1 integrin in cells treated with siCTRL or siTLN1 #3 and siTLN2 #3 oligos (siCTRL, n = 799 filopodia; siTLN1 #3 and siTLN2 #3, n = 889 filopodia; three biological repeats). For all panels, p-values were determined using a randomization test. NS indicates no statistical difference between the mean values of the highlighted condition and the control.

Talin is a critical regulator of integrin activity, known to localize to and modulate filopodia function (Lagarrigue et al., 2015; Jacquemet et al., 2016), and has been predicted by us and others to trigger integrin activation at filopodia tips (Jacquemet et al., 2019; Lagarrigue et al., 2015). To validate this notion, cells silenced for TLN1 and TLN2 were plated on fibronectin and stained for active β1 integrin (Fig. 2B). Given that full silencing of both talin isoforms render cells poorly adherent (Zhang et al., 2008), we aimed at partial silencing to ensure sufficient cell adhesion to the coverslip (TLN1 60% silencing efficiency, TLN2 100% silencing efficiency; Fig. S1I). Reduced talin expression did not affect filopodia length (Fig. 2C) but was sufficient to clearly decrease active β1 integrin localization at filopodia tips as well as the percentage of filopodia containing active β1 integrin at their tips (Fig. 2D and 2E). Altogether, our data demonstrate that talin is required for integrin activation at filopodia tips.

### The FERM domain of MYO10 is required for integrin activation but not localization at filopodia tips

We previously observed that FMNL3-induced filopodia rarely contain active β1 integrin (Jacquemet et al., 2019). A careful reanalysis of these data, using intensity profile mapping, indicates that active β1 integrin can be detected in only 23 % of FMNL3-induced filopodia (Fig. S3). However, this is not due to an absence of β1 integrin as all FMNL3-induced filopodia are strongly positive for inactive β1 integrins (Fig. S3). As integrin activation is a prominent feature of MYO10-positive filopodia (Fig. 1), we hypothesized that MYO10 could functionally contribute to integrin activation in filopodia tips.

MYO10 directly binds to integrins via its FERM domain (Hirano et al., 2011; Zhang et al., 2004). In this context, MYO10 is thought to actively transport integrins as well as other cargo to filopodia tips. We assessed the contribution of the MYO10 FERM domain to integrin localization in filopodia by creating a FERM domain deletion construct (MYO10^ΔF^) (Fig. 3A). We carefully designed this construct by taking into consideration the previously reported MYO10-FERM domain structures (PDB ids: 3PZD and 3AU5) (Wei et al., 2011; Hirano et al., 2011). MYO10^ΔF^ was expressed in U2-OS cells, which express low levels of endogenous MYO10 (Young et al., 2018; Jacquemet et al., 2016). Deletion of the MYO10-FERM domain led to a small but significant reduction in filopodia number and filopodia length, in line with previous reports (Zhang et al., 2004; Watanabe et al., 2010) (Fig. 3B-D). Strikingly, the majority of MYO10^ΔF^-filopodia (80%) were devoid of active β1 integrins at their tips (Fig. 3E-G) while the uniform distribution of inactive β1 integrins along the filopodium length remained unaffected (Fig. 3E-G). In line with these results, MYO10^ΔF^-induced filopodia were unstable and unable to attach to the underlying ECM (Fig. 3H and Video 1). Taken together, these findings demonstrate that MYO10-FERM is required for integrin activation at filopodia tips but not for β1 integrin localization to filopodia tips (Fig. 3 and Fig. S2).

**Figure 3.**
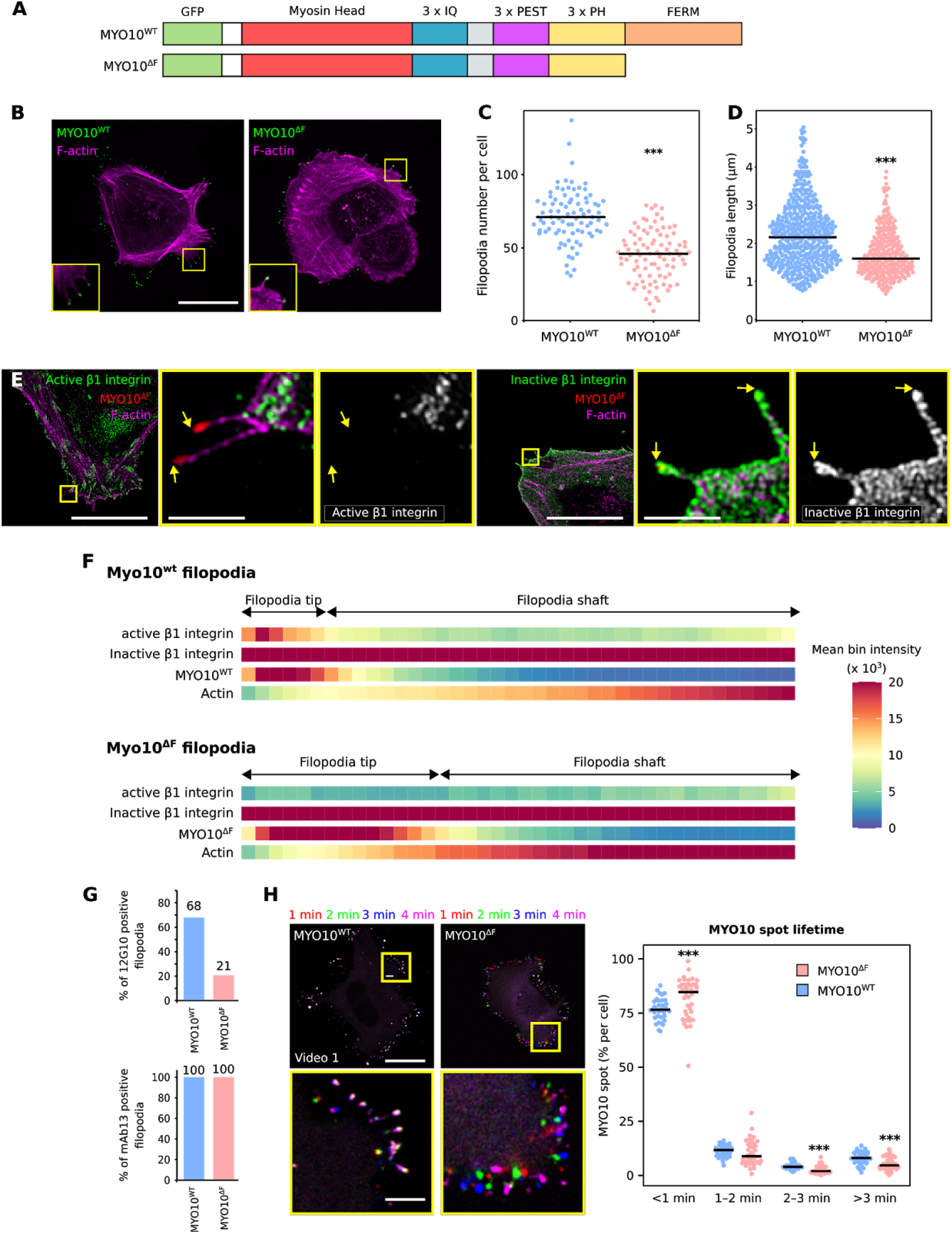
MYO10-FERM is required for integrin activation in filopodia. **(A**) Cartoon illustrating the EGFP-MYO10^WT^ and EGFP-MYO10^ΔF^ constructs. **(B-C**) U2-OS cells transiently expressing EGFP-MYO10^WT^ or EGFP-MYO10^ΔF^ were plated on fibronectin for 2 h, fixed, and imaged using a spinning-disk confocal microscope. (**B**) Representative MIPs are displayed. Scale bar: 25 μm. (**C**) The number of MYO10-positive filopodia per cell was then quantified (n > 85 cells, three biological repeats; *** p-value <0.001). **(D**) Quantification of MYO10^WT^ and MYO10^ΔF^ filopodia length from SIM images (EGFP-MYO10^WT^, n = 623 filopodia; EGFP-MYO10^ΔF^, n = 283 filopodia; three biological repeats; *** p value = <0.001). **(E**) U2-OS cells expressing EGFP-MYO10^ΔF^ were plated on fibronectin for 2 h, stained for active (antibody 12G10) or inactive (antibody mAb13) β1 integrin and F-actin, and imaged using SIM. Representative MIPs are displayed. The yellow squares highlight ROIs, which are magnified; yellow arrows highlight filopodia tips; scale bars: (main) 20 μm; (inset) 2 μm. **(F**) Heatmap highlighting the sub-filopodial localisation of the proteins stained in (**E**) generated from their intensity profiles (EGFP-MYO10^WT^: MYO10, n = 623 filopodia; F-actin, n = 623 filopodia; active β1 integrin, n = 329 filopodia; inactive β1 integrin, n = 369 filopodia. EGFP-MYO10^ΔF^: MYO10, n = 283 filopodia; F-actin, n = 283 filopodia; active β1 integrin, n = 347 filopodia; inactive β1 integrin, n = 250 filopodia. Three biological repeats). **(G**) Bar chart highlighting the percentage of MYO10^WT^ and MYO10^ΔF^-induced filopodia with detectable levels of active (12G10) and inactive (mAb13) β1 integrin (12G10: MYO10^WT^, n = 329 filopodia; MYO10^ΔF^, n = 347 filopodia; mAb13: MYO10^WT^, n = 369 filopodia; MYO10^ΔF^, n = 250 filopodia; three biological repeats). **(H**) U2-OS cells expressing EGP-MYO10^WT^ or EGFP-MYO10^ΔF^ were plated on fibronectin and imaged live using an Airyscan confocal microscope (1 picture every 5 s over 20 min; scale bar = 25 μm; Video 1). For each condition, MYO10-positive particles were automatically tracked, and MYO10 spot lifetime (calculated as a percentage of the total number of filopodia generated per cell) was plotted and displayed as boxplots (see Methods for details; three biological repeats; EGP-MYO10^WT^, n = 33 cells; EGFP-MYO10^ΔF^, n = 38 cells, *** p-value < 0.006). For all panels, p-values were determined using a randomization test.

As these findings challenge the model of the MYO10 FERM domain acting as a cargo-transporter of integrin to filopodia tips, we tested whether the presence of inactive β1 integrins in MYO10^ΔF^-filopodia could be due to the low endogenous MYO10 present in these cells. We expressed wild-type or MYO10^ΔF^ in MYO10-silenced U2-OS cells (90% silencing efficiency with a 3’ UTR-targeting RNA oligo) and analyzed β1 integrin distribution using SIM (Fig. S3A). Inactive β1 integrin localization in MYO10^ΔF^-filopodia was not affected by the silencing of endogenous MYO10, further validating that MYO10-FERM is not required to localize β1 integrin to filopodia (Fig. S3B-D). Interestingly, silencing of endogenous MYO10 led to a small decrease in the percentage of MYO10 filopodia that contain active integrin at their tips, suggesting that integrin activation at filopodia tips by MYO10 may be dose-dependent (Fig. S3D).

### MYO10-FERM deletion does not influence the localization of established filopodia tip components

As MYO10-FERM is thought to be the major cargo binding site in MYO10 (Wei et al., 2011), we hypothesized that the lack of integrin activation at the tip of MYO10^ΔF^ filopodia would be due to the absence of a key integrin activity modulator. We co-expressed six established filopodia tip components (Jacquemet et al., 2019), TLN1, FERMT2, CRK, DIAPH3, BCAR1, and VASP, with either MYO10^WT^ or MYO10^ΔF^. SIM microscopy revealed that the localization of these proteins was unaffected by MYO10-FERM domain deletion (Fig. S4). Interestingly, VASP has been previously described as an MYO10-FERM cargo but its localization at filopodia tips was clearly unaffected by MYO10-FERM deletion (Young et al., 2018; Tokuo and Ikebe, 2004; Lin et al., 2013). Altogether, our results demonstrate that the recruitment of key filopodia tip proteins, including TLN1, is independent of the MYO10 FERM domain and suggest that MYO10-FERM may regulate integrin activity via another mechanism than cargo transport.

### The interaction between MYO10 and integrins regulates integrin activation at filopodia tips

MYO10-FERM is composed of four subdomains, namely a MyTH subdomain and three FERM lobes F1, F2, and F3. To further dissect which part of MYO10-FERM is responsible for mediating integrin activation at filopodia tips, two additional MYO10 deletion constructs were generated where either the F2F3 (MYO10^ΔF2F3^) or the F3 (MYO10^ΔF3^) lobes are missing (Fig. S5A). We expressed MYO10^ΔF2F3^, MYO10^ΔF3^, MYO10^ΔF^ and MYO10^WT^ in U2-OS cells and compared their filopodia properties (Fig. S5B-E). MYO10^ΔF2F3^ and MYO10^ΔF3^ filopodia were shorter than MYO10^WT^ filopodia but longer than MYO10^ΔF^ filopodia indicating that the MyTH, F1 and F3 subdomains contribute to filopodia elongation (Fig. S5C). Importantly, MYO10^ΔF2F3^ and MYO10^ΔF3^ filopodia displayed low amounts of active β1 integrin at their tips indicating that the MYO10 F3 subdomain is required to activate integrin at filopodia tips (Fig. S5D-E). As others have shown that the MYO10 F3 subdomain contains the integrin β1 binding site (Zhang et al., 2004), our results led us to speculate that MYO10 needs to interact with integrin directly to promote integrin activation.

While the site where β1 integrin binds to MYO10-FERM remains unknown, the integrin-binding site has been mapped in talin-FERM. Despite some controversy regarding the full talin-FERM structure, superimposition of talin and MYO10 FERM domains revealed that both adopt a similar fold in the β integrin tail binding subdomains (Fig. 4A and FigS6A) (Zhang et al., 2020; Elliott et al., 2010). Therefore, we can predict mutations likely to disturb the MYO10-integrin interaction (S2001_F2002insA and T2009D, Fig. 4B). The introduction of these mutations in MYO10-FERM (FERM^ITGBD^) led to a 64 % reduction in the ability of β1 integrin tail peptides to pulldown GFP-tagged MYO10-FERM domains from cell lysate indicating that these mutations can impede the interaction between MYO10 and integrins (Fig. 4C). Cells expressing full-length MYO10^ITGBD^ generated filopodia to the same extent as cells expressing MYO10^WT^ (Fig. 4D and 4E), but MYO10^ITGBD^ filopodia were shorter than MYO10^WT^ filopodia (Fig. 4F). Notably, only 25 % of MYO10^ITGBD^ filopodia contained detectable levels of active β1 integrin at their tips (Fig. 4G and 4H). Thus, we conclude that an intact integrin-binding site within MYO10-FERM is required for MYO10 to fully activate β1 integrin at filopodia tips.

**Figure 4.**
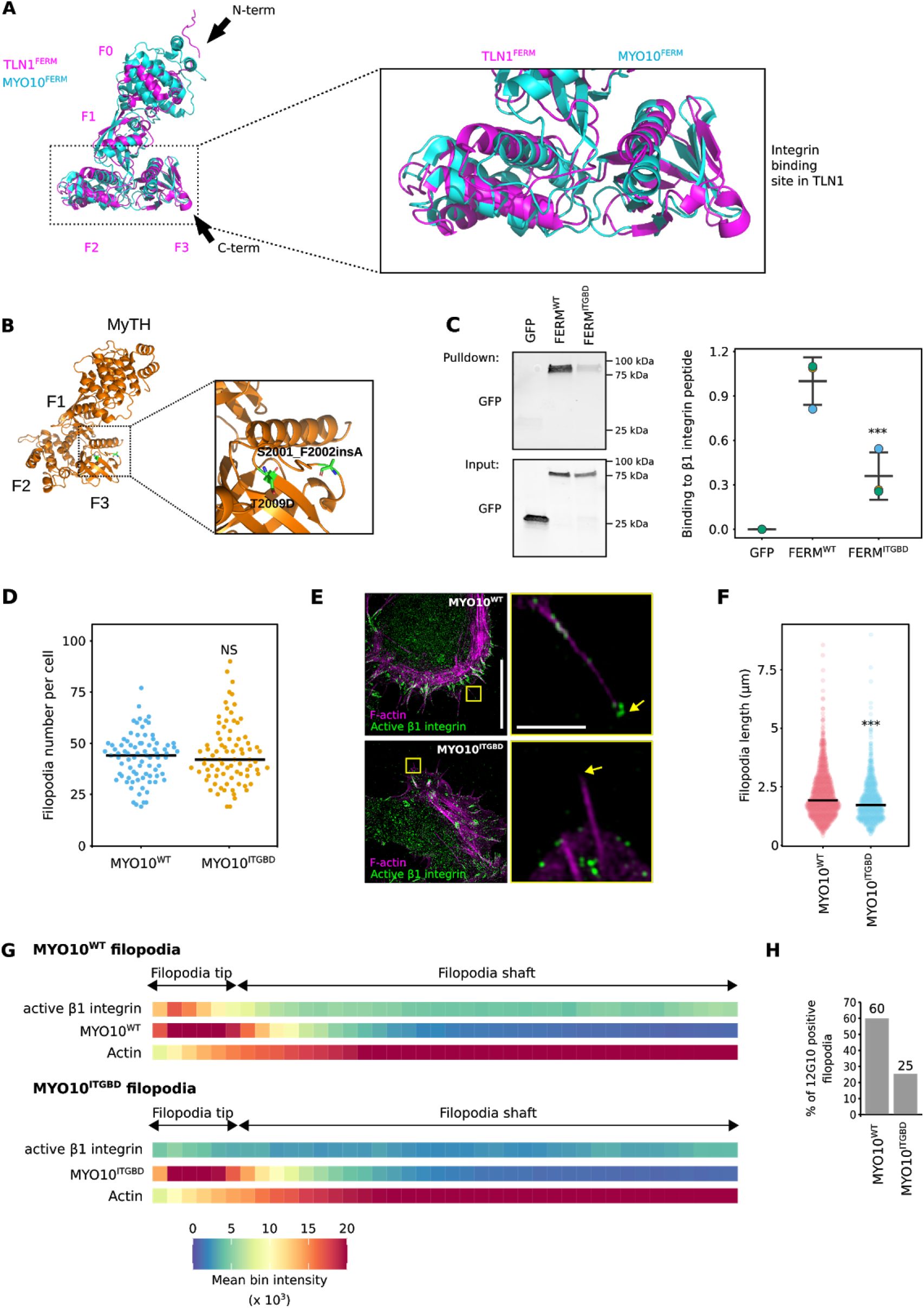
An intact integrin-binding site within MYO10-FERM is required for MYO10-mediated integrin activation at filopodia tips. **(A**) Visualisation of the structure of MYO10-FERM (PDB: 3PZD; (Wei et al., 2011)) and TLN1-FERM (PDB: 6VGU (Zhang et al., 2020), domains using PyMOL. The black arrows indicate the protein orientation from N to C terminal. The two FERM domains were superimposed to highlight their structural homology and differences. The integrin-binding region on the talin-FERM domain is highlighted and magnified. **(B**) The structure of the MYO10-FERM mutated on the predicted integrin-binding site was modeled using SWISS-MODEL (Waterhouse et al., 2018) based on the known MYO10-FERM wild-type structure (PDB: 3PZD) and was visualized using PyMOL. The mutated residues are highlighted in green. (**C**) β1-integrin tail peptide pulldown in U2-OS cells expressing EGFP-tagged MYO10-FERM WT (FERM^WT^) or mutant (FERM^ITGBD^) or EGFP alone. MYO10-FERM recruitment to the β1-integrin tail was assessed using western blot (n = 3, *** p-value = 0.008 and was calculated using a Welch’s t-test). A representative western blot, as well as the western blot quantifications, are displayed. Individual repeats are color-coded (Lord et al., 2020; Goedhart, 2020). **(D**) U2-OS cells transiently expressing full-length EGFP-MYO10^WT^ or EGFP-MYO10^ITGBD^ were plated on fibronectin for 2 h, fixed and imaged using a spinning-disk confocal microscope, and the number of MYO10-positive filopodia per cell was quantified (EGP-MYO10^WT^, n = 81 cells; EGFP-MYO10^ITGBD^, n = 81 cells; three biological repeats). The p-value was determined using a randomization test. NS indicates no statistical difference between the mean values of the highlighted condition and the control. **(E-H**) U2-OS cells expressing EGFP-MYO10^WT^ or EGFP-MYO10^ITGBD^ were plated on fibronectin for 2 h, stained for active β1 integrin (antibody 12G10) and F-actin, and imaged using SIM. (**E**) Representative MIPs are displayed. The yellow squares highlight ROIs, which are magnified; yellow arrows highlight filopodia tips; scale bars: (main) 20 μm; (inset) 2 μm. (**F**) Quantification of MYO10^WT^ and MYO10^ITGBD^ filopodia length from SIM images (MYO10^WT^, n = 1073 filopodia; MYO10^ITGBD^ n = 693 filopodia ; three biological repeats; *** p-value = <0.001). The p-value was determined using a randomization test. (**G**) Heatmap highlighting the sub-filopodial localization of the indicated proteins based on their intensity profiles. (EGFP-MYO10^WT^, n = 1073 filopodia; EGFP-MYO10^ITGBD^, n = 693 filopodia; three biological repeats). (**H**) Bar chart highlighting the percentage of MYO10^WT^ and MYO10^ITGBD^ filopodia with detectable levels of active β1 integrin (MYO10^WT^, n = 1073 filopodia; MYO10^ITGBD^ n = 693 filopodia; three biological repeats).

### Unlike Talin-FERM, MYO10-FERM domain alone is not able to activate integrins

The talin-FERM domain is necessary and sufficient to activate integrins (Anthis et al., 2009; Lilja et al., 2017). Given our data indicating that MYO10-FERM is required to activate integrin at filopodia tips (Fig 3 and 4), we tested whether MYO10-FERM could modulate integrin activity similarly to talin-FERM. We employed a flow cytometric assay to measure active cell-surface integrins relative to total cell-surface integrins (Lilja et al., 2017) (Fig. 5A-C). As expected overexpression of the talin-FERM domain significantly increased integrin activity (Fig. 5A). In contrast, overexpression of the MYO10-FERM domain failed to activate integrins and instead led to a small but highly reproducible decrease in integrin activity in CHO and U2-OS cells (Fig. 5A-B). Similar data were obtained in U2-OS cells overexpressing full-length MYO10 (Fig. 5B). Conversely, silencing of MYO10 increased integrin activity in MDA-MB-231 cells, where mutant p53 drives high endogenous MYO10 levels (Arjonen et al., 2014), and this was reversed by reintroduction of full-length MYO10 (Fig. 5C and S6B). Consistent with decreased integrin activation, MYO10-FERM expression attenuated cell adhesion/spreading on fibronectin over time as measured with the xCELLigence apparatus (cell spreading assay system based on electrical impedance) or by measuring cell spreading area (Fig. 5D-F) (Hamidi et al., 2017). Altogether our data indicate that, even though the MYO10 FERM domain is necessary for spatially restricted integrin activation at filopodia tips, the MYO10-FERM domain alone is not capable of activating integrins.

**Figure 5.**
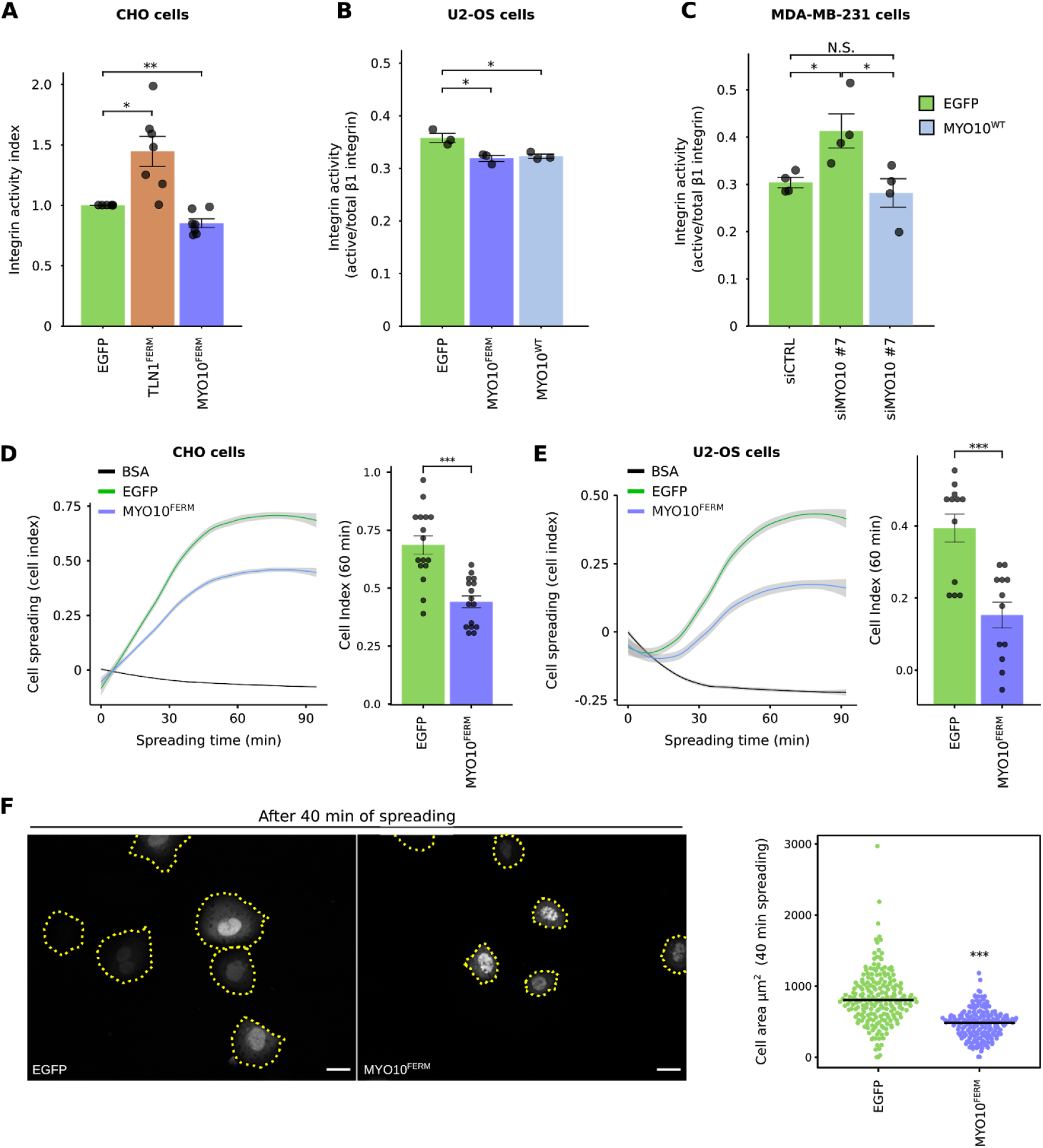
The MYO10 FERM domain inhibits integrin activity. **(A**) CHO cells expressing EGFP or the EGFP-tagged FERM domains of TLN1 (EGFP-TLN1^FERM^) or MYO10 (EGFP-MYO10^FERM^) were either incubated with an Alexa647-labeled fibronectin fragment (FN7–10) and fixed, or fixed directly and stained with a (total) anti-ITGA5 antibody (anti-hamster VLA5 antibody PB1). Samples were then analyzed by FACS. The integrin activity index was calculated using the fibronectin and the ITGA5 signals as described in the Methods. Results are displayed as bar charts where the individual experiments are highlighted as dots, and error bars represent the standard error of the mean (** p-value = 0.0062, n = 7 of biological repeats). The p-value was determined using a one-sample t-test. (missing stats: EGFP vs TalinFERM p = 0.012 one-sample t-test. TalinFERM vs Myo10FERM p = 0.003 two-sample t-test). **(B**) U2-OS transiently expressing various EGFP constructs (as indicated), were fixed and stained for active (antibody 9EG7) or total β1 integrin (antibody P5D2). Staining intensity was recorded by flow cytometry and integrin activation was assessed as a ratio between active and total integrin (9EG7/P5D2 ratio). Error bars indicate standard error of the mean (* p-value < 0.05; C, n = 5 biological repeats; D, n = 4 biological repeats). The p-values were determined using a Student’s two-tailed t-test. **(C**) MDA-MB-231 cells, previously silenced for MYO10 (siMYO10 #7 oligo targets the 3’ UTR of MYO10 mRNA) and expressing EGFP or EGFP-MYO10, were fixed and stained for active (antibody 9EG7) or total β1 integrin (antibody P5D2). Staining intensity was recorded by flow cytometry and integrin activation was assessed as a ratio between active and total integrin (9EG7/P5D2 ratio). Error bars indicate standard error of the mean (* p-value < 0.05; n = 4 biological repeats). The p-value was determined using a Student’s two-tailed t-test. **(D-E**) CHO or U2-OS cells transiently expressing EGFP or EGFP-MYO10^FERM^ were left to adhere on fibronectin and their spreading was monitored over time using the xCELLigence system. The cell index over time is displayed, grey areas indicate the 95% confidence intervals. The cell index at 60 min is also displayed as a bar chart (error bars indicate the standard error of the mean, *** p-value < 0.001, n = 4 (CHO) and 3 (U2-OS) biological repeats). The p-value in bar charts was determined using a Student’s two-tailed t-test. **(F)** U2-OS cells transiently expressing EGFP or EGFP-MYO10^FERM^ were seeded on fibronectin and allowed to spread for 40 min prior to fixation. Samples were imaged using a confocal microscope and the cell area measured using Fiji (*** p-value < 0.001; EGFP, 208 cells; EGFP-MYO10^FERM^, 188 cells; n = 3 biological repeats). The p-values were determined using a randomization test. Scale bars are 16 μm.

### Unlike Talin-FERM, MYO10-FERM binds to both α and β integrin tails

Despite being homologous domains with high structural similarity, the functional difference between MYO10 FERM and Talin FERM domains in their ability to activate integrins prompted us to compare their binding affinities to integrin cytoplasmic tails. Recombinant His-tagged MYO10 and talin-FERM were expressed in bacteria, purified (Fig S6C), and their binding affinity to integrin α and β tails was measured using microscale thermophoresis (Fig. 6A and Fig. 6B; see methods for details) (Jerabek-Willemsen et al., 2014). As expected, talin-FERM interacted with the β1 integrin tail (measured affinity of 4.7 μM) but not to α integrin tails (Goult et al., 2009). This result is in agreement with measurements done by others using the same method (Haage et al., 2018). Interestingly MYO10-FERM bound to the β1 integrin tail with slightly lower affinity than talin-FERM (measured affinity of 25.1 μM) (Fig 6A and 6B). This result indicates that talin may be able to outcompete MYO10 for integrin binding.

**Figure 6.**
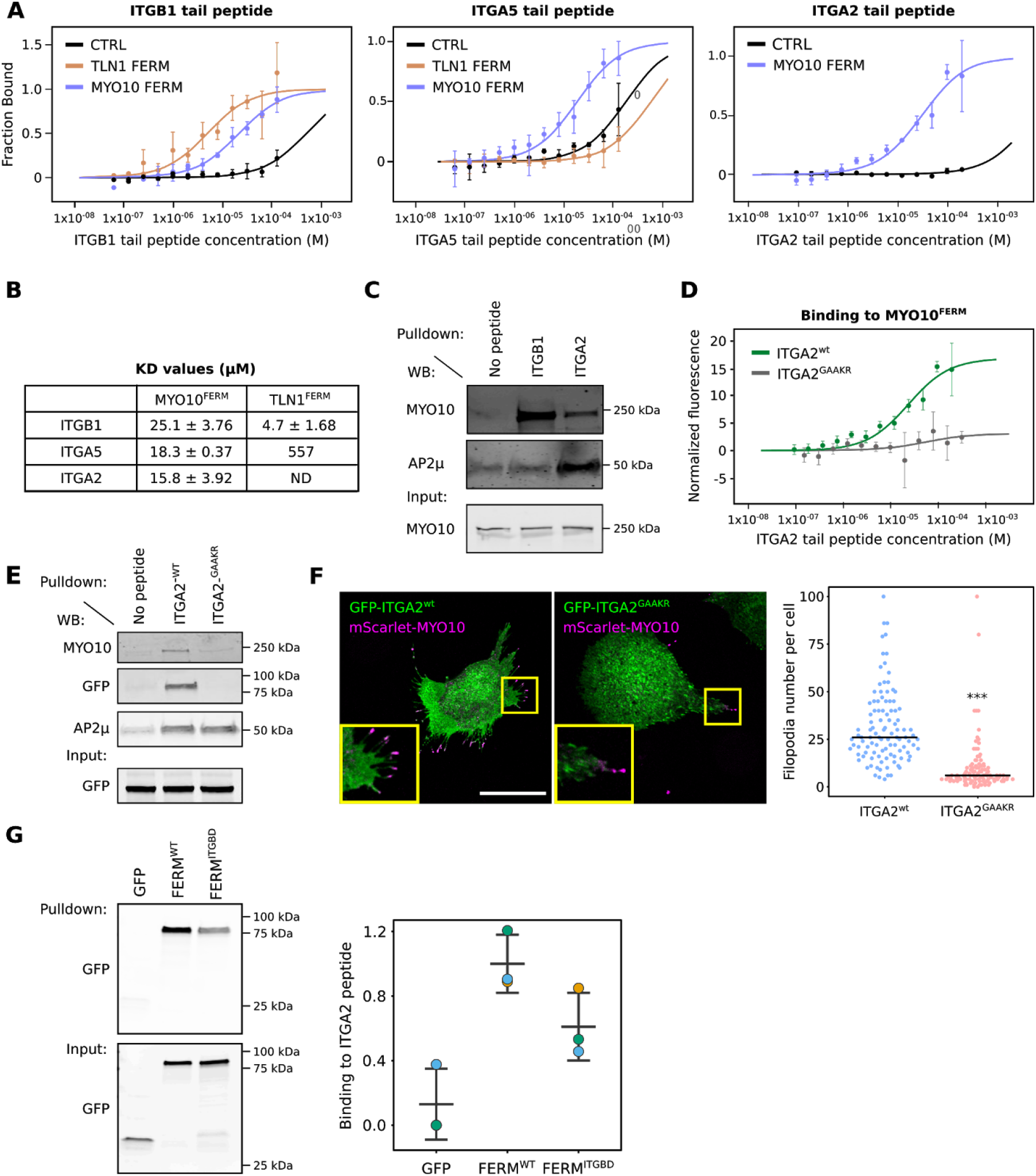
MYO10 binds to both α and β integrin tails. **(A**) Recombinant 6xHis-tagged FERM domains from TLN1 (TLN1^FERM^) and MYO10 (MYO10^FERM^) and a 6xHis peptide (CTRL) were labeled and their binding to integrin tails was recorded using microscale thermophoresis. In these experiments, 20 nM of labeled FERM was used while the integrin tail peptides were used at increasing concentrations. Graphs were generated by pooling together 3 independent experiments (see Methods for details). **(B**) Table showing the Kd values obtained when measuring the binding, using microscale thermophoresis, of TLN1^FERM^ and MYO10^FERM^ to the indicated integrin tail peptides. For each condition, data from 3 independent experiments were pooled together to obtain Kd values. **(C**) Integrin tail pull-downs were performed from U2-OS cell lysates using magnetic beads. The recruitment of MYO10 and AP2μ was then analyzed by western blot (n = 3 biological experiments). A representative western blot is displayed. **(D**) Recombinant MYO10-FERM domain (MYO10^FERM^) was labeled and its binding to the intracellular tails of wild-type ITGA2 (ITGA2^WT^) or of ITGA2 mutated on the GFFKR consensus site (ITGA2^GAAKR^) was recorded using microscale thermophoresis. In these experiments, 20 nM of MYO10-FERM was used while the integrin tail peptides were used at increasing concentration. Graphs were generated by pooling together 3 independent experiments (see methods for details). **(E**) Integrin tail pull-downs were performed from cell lysate generated from U2-OS cells stably expressing EGFP-MYO10^FERM^. The recruitment of endogenous MYO10, EGFP-MYO10^FERM^, and AP2μ was then analyzed by western blot (n = 3 biological experiments). A representative western blot is displayed. **(F**) CHO cells transiently expressing mScarlet-MYO10 and full-length GFP-ITGA2^WT^ or GFP-ITGA2^GAAKR^ were plated on collagen I for 2 h, fixed, and imaged using a spinning-disk confocal microscope. Representative MIPs are displayed. Scale bar: 25 μm. The number of MYO10-positive filopodia per cell was then quantified (n > 107 cells, four biological repeats; *** p-value < 0.001). P-values were determined using a randomization test. (**G**) Different EGFP-tagged MYO10 FERM domains or EGFP alone were pulled down from U2-OS lysate using α2-integrin tail peptide coupled beads. MYO10 FERM recruitment to α2-integrin tail was assessed using western blot (n = 3 biological experiments). A representative western blot as well as the western blot quantifications are displayed. Individual repeats are color-coded (Lord et al., 2020; Goedhart, 2020).

Unexpectedly, our results indicated that, in contrast to talin-FERM, α integrin-tails also interact with MYO10-FERM in vitro (Fig. 6A and Fig. 6B) and with endogenous MYO10 in cell lysate (Fig. 6C). The ability of MYO10 to interact with both α- and β-tail peptides appeared to be specific as the clathrin adaptor AP2μ, a known α2 integrin tail specific binder (De Franceschi et al., 2016), was pulled down only with the α2 integrin tail (Fig. 6C). The MYO10-α-tail interaction was dependent on the highly conserved membrane-proximal GFFKR motif, present in most integrin α-tails (De Franceschi et al., 2016). Mutation of the motif in the α2-integrin tail (FF/AA mutation, named ITGA2^GAAKR^) abolished the binding of recombinant MYO10-FERM in vitro (Fig. 6D), and in pull-down with full-length MYO10 (Fig. 6E). Importantly, AP2μ recruitment was unaffected by the mutation (AP2μ binds to a separate motif in the α2-tail) (Fig. 6E). Together, these experiments demonstrate that MYO10 binds to integrin β-tails, in line with previous reports (Zhang et al., 2004; Hirano et al., 2011), and reveal a previously unknown interaction between MYO10-FERM and the GFFKR motif in integrin α tails. Binding to both integrin tails has been demonstrated as a mechanism for FLNA-mediated integrin inactivation (Liu et al., 2015) and, thus, may be the underlying reason for the inability of MYO10-FERM to activate integrins.

To test the relevance of the GFFKR α-integrin tail motif in filopodia induction, we expressed full-length wild-type ITGA2 and ITGA2^GAAKR^ in CHO cells (these cells lack endogenous collagen-binding integrins) and investigated MYO10 filopodia formation on collagen I (Fig. 6F). ITGA2^GAAKR^ localizes to the plasma membrane and is expressed at similar levels to wild-type in CHO cells (Alanko et al., 2015). ITGA2^GAAKR^-expressing cells generated less filopodia than cells expressing wild-type ITGA2 indicating that the GFFKR motif in the ITGA2 tail contributes to filopodia formation. We could not, however, directly assess the relevance of the MYO10-α integrin interaction to filopodia functions as the MYO10^ITGBD^ construct also displayed reduced binding toward ITGA2 (Fig. 6G).

### MYO10-FERM domain fine-tunes integrin activity at filopodia tips

To further investigate how MYO10-FERM regulates integrin activity in filopodia and the functional differences between talin and MYO10 FERM domains, we created a chimera construct, where the FERM domain from MYO10 was replaced by the one from TLN1 (MYO10^TF^) (Fig. 7A). Both MYO10^WT^, MYO10^TF^ strongly accumulated at filopodia tips (Fig. 7B and 7C). Interestingly, in a small proportion of cells (below 1%), MYO10^TF^ also localized to enlarged structures connected to stress fibers that are reminiscent of focal adhesions (Fig. 7C).

**Figure 7.**
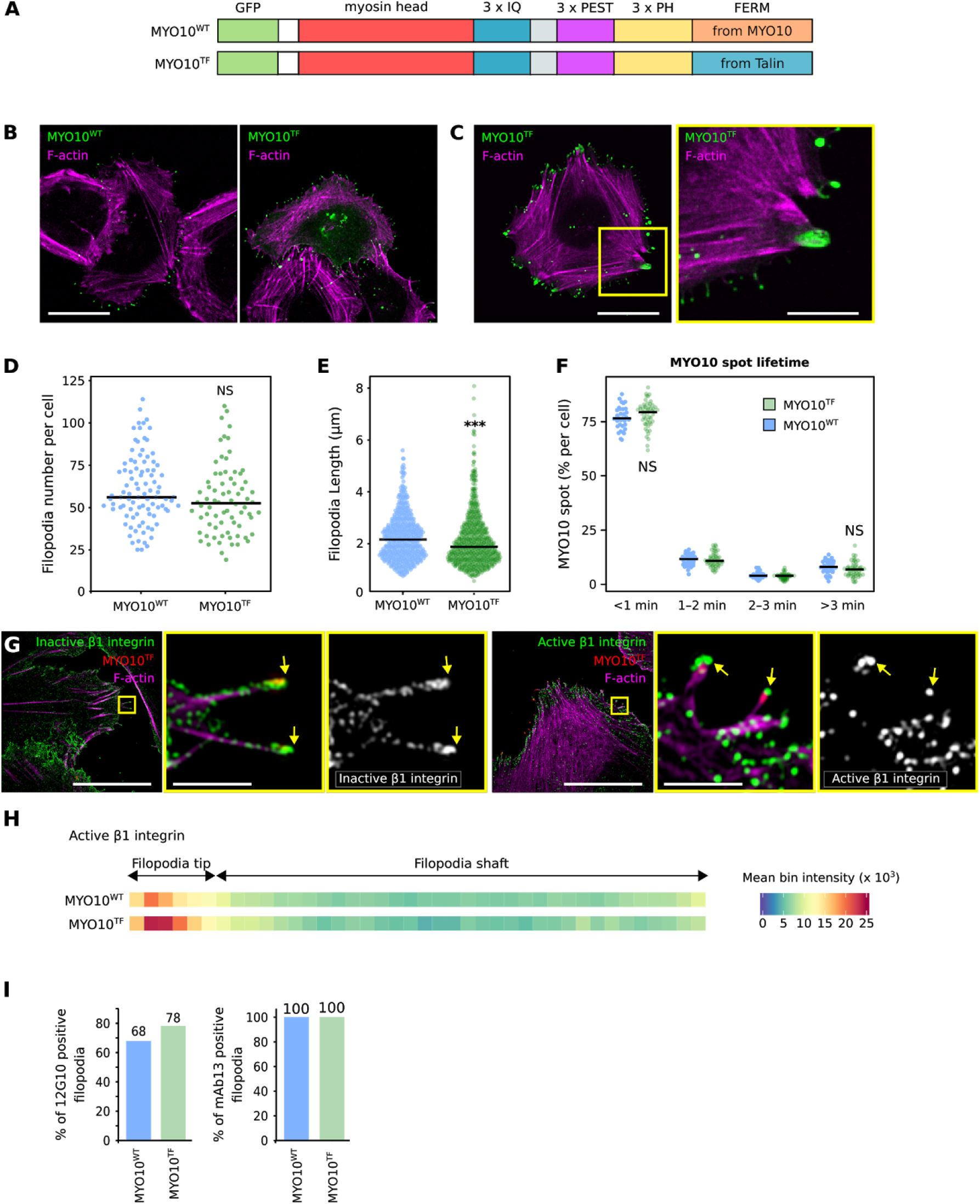
MYO10-FERM fine-tunes integrin activity at filopodia tips. **(A**) Cartoon illustrating the EGFP-MYO10^WT^ and EGFP-MYO10^TF^ constructs. **(B-E**) U2-OS cells transiently expressing EGFP-MYO10^WT^ or EGFP-MYO10^TF^ were plated on fibronectin for 2 h, fixed and imaged using a spinning-disk confocal microscope or an Airyscan confocal microscope. **B**) Representative MIPs acquired on a spinning-disk confocal are displayed. Scale bar: 25 μm. **C**) An image acquired on an Airyscan confocal microscope is displayed. The yellow square highlights an ROI, which is magnified. scale bars: (main) 25 μm; (inset) 5 μm. **(D**) The number of MYO10-positive filopodia per cell was quantified (EGP-MYO10^WT^, n = 93 cells; EGFP-MYO10^TF^, n = 74 cells; three biological repeats). **(E**) Quantification of MYO10^WT^ and MYO10^TF^ filopodia length from SIM images (EGFP-MYO10^WT^, n = 512 filopodia; EGFP-MYO10^TF^ n = 669 filopodia ; three biological repeats; *** p-value = <0.001). **(F**) U2-OS cells expressing EGFP-MYO10^WT^ or EGFP-MYO10^TF^ were plated on fibronectin and imaged live using an Airyscan confocal microscope (1 picture every 5 s over 20 min). For each condition, MYO10-positive particles were automatically tracked, and MYO10 spot lifetime (calculated as a percentage of the total number of filopodia generated per cell) was plotted and displayed as boxplots (three biological repeats, EGP-MYO10^WT^, n= 33 cells; EGFP-MYO10^TF^, n= 53 cells). **(G**) U2-OS cells expressing EGFP-MYO10^TF^ were plated on fibronectin for 2 h, stained for active (antibody 12G10) or inactive (antibody mAb13) β1 integrin and F-actin, and imaged using SIM. Representative MIPs are displayed. The yellow squares highlight ROIs, which are magnified; yellow arrows highlight filopodia tips; scale bars: (main) 20 μm; (inset) 2 μm. **(H**) Heatmap highlighting the sub-filopodial localization of active β1 integrin in cells expressing EGFP-MYO10^WT^ or EGFP-MYO10^TF^ (EGFP-MYO10^WT^, n = 512 filopodia; EGFP-MYO10^TF^, n = 669 filopodia; three biological repeats). **(I**) Bar chart highlighting the percentage of MYO10^WT^ and MYO10^TF^ filopodia with detectable levels of active and inactive β1 integrin (active β1 integrin: MYO10^WT^, n = 329 filopodia; MYO10^TF^ n = 255 filopodia; inactive β1 integrin: MYO10^WT^, n = 369 filopodia; MYO10^TF^ n = 414 filopodia; three biological repeats).

Cells expressing MYO10^TF^ generated filopodia to the same extent as cells expressing MYO10^WT^ (Fig. 7D). MYO10^TF^ filopodia were slightly shorter than MYO10^WT^ filopodia but of comparable dynamics (Fig. 7E-F). These results show that the talin-FERM can replace the MYO10-FERM domain and highlight an unanticipated level of interchangeability between integrin-binding FERM domains in regulating filopodia properties. Importantly, active β1 integrin accumulated more efficiently at the tips of MYO10^TF^ filopodia (Fig. 7G and 7H) and MYO10^TF^ filopodia were more likely to contain active β1 integrin at their tips than MYO10^WT^ filopodia (Fig. 7G-I). Silencing of TLN1 and TLN2 still impeded MYO10^TF^ filopodia formation, indicating that talin-FERM fused to the MYO10 motor is insufficient to substitute for the lack of endogenous full-length talin (Fig. S6C and D). The increased amount of active β1 integrin at the tip of MYO10^TF^ filopodia is likely due to the ability of talin-FERM to activate integrin directly (Fig. 5) and/or because talin-FERM binds to integrins with a higher affinity than MYO10-FERM (Fig. 6). Altogether, our data indicate that an integrin-binding proficient FERM domain coupled to a myosin motor is required to activate, but not to transport, integrin in filopodia (Fig. 2 and 5). We also found that the nature of this FERM domain, whether it is capable of activating or inactivating integrin, contributes to fine-tuning integrin activation at filopodia tips (Fig. 5).

## Discussion

Here, we observed that active integrin accumulates at filopodia tips while inactive integrin localizes throughout filopodia shafts. We find that integrin activation in filopodia is uncoupled from focal adhesions or the actomyosin machinery but is instead regulated by talin and MYO10. Contrary to previous assumptions, MYO10 is not required to localize integrin to filopodia, but its integrin-binding FERM domain is required for integrin activation at filopodia tips. We find, however, that unlike talin-FERM, MYO10-FERM itself does not promote integrin activation. Swapping MYO10-FERM with talin-FERM, in the context of full-length MYO10, and mutation of the integrin-binding site in MYO10 demonstrate that an integrin-binding FERM domain coupled to a myosin motor is a core requirement for integrin activation at filopodia tips.

We find that MYO10-FERM interaction with integrins is required to localize active integrin to filopodia tips. The simplest assumption would be that MYO10, in its typical capacity as a myosin motor, specifically transports active integrin to filopodia tips. However, our data suggest otherwise as 1) the MYO10 FERM domain alone inactivates integrins and therefore integrins would not be in an active state during transport, 2) talin is required to localize active integrins at filopodia tips, and 3) integrin activation is thought to be a fast and tightly regulated process (Sun et al., 2019), all evidence pointing to an on-site integrin activation mechanism in filopodia tips. In addition, direct transport of integrin by MYO10 to filopodia tips has yet to be formally observed. Instead, we propose that integrins localize along the filopodia plasma membrane via membrane diffusion and are activated at filopodia tips in a two-step process by MYO10 and talin. In this model, MYO10 could tether integrins at filopodia tips due to its motor domain and provide resistance against the actin retrograde flow present in filopodia (Bornschlögl et al., 2013) allowing sufficient time for talin-mediated activation.

A role for talin in mediating integrin activity at filopodia tips is not surprising and was predicted by us and by others (Lagarrigue et al., 2015; Jacquemet et al., 2016). We have demonstrated using live-cell imaging that, in filopodia, talin and MYO10 always co-localize (Jacquemet et al., 2019), indicating that talin-mediated integrin activation is not modulated by talin recruitment. As the small GTPase Rap1 localizes to filopodia and is required to support filopodia functions (Jacquemet et al., 2016; Lagarrigue et al., 2015), a plausible mechanism would be that, upon filopodia initiation, talin is kept in an auto-inhibited conformation at filopodia tips. Once activated by Rap1 (directly or indirectly via RIAM), talin auto-inhibition is released, and talin associates with and activates integrins, triggering adhesion (Sun et al., 2019). In this context, Rap1 could be activated by both intracellular and extracellular signals, including calcium entry via calcium channels (Efremov et al., 2020; Jacquemet et al., 2016).

The precise mechanisms favoring integrin binding to MYO10 or talin in filopodia remain to be elucidated. One possibility is that talin-FERM outcompetes MYO10-FERM. Indeed, our in vitro experiments indicate that talin-FERM has, in solution, a higher affinity for integrin β-tail compared to MYO10-FERM. In addition, talin affinity for β integrin tails will be even stronger in cells due to the presence of negatively charged membrane phosphoinositides that interact with talin-FERM (Chinthalapudi et al., 2018; Franceschi et al., 2019), and which are known to accumulate at filopodia tips (Jacquemet et al., 2019). Interestingly, while MYO10 and Talin FERM domains structurally adopt a very similar fold, we find that these two FERM domains are functionally distinct. MYO10-FERM is not capable of directly activating integrin and can interact with both integrin tails. Yet, remarkably, swapping MYO10-FERM with talin-FERM fully supported filopodia function and integrin activation at filopodia tips, suggesting an unanticipated interchangeability between these FERM domains in spatially regulating integrin activation in filopodia. As other FERM domain-containing myosins, including MYO7 and MYO15, also localize to filopodia tips (Jacquemet et al., 2019; Arthur et al., 2019), where their roles are mostly unknown, future work will examine the contribution of these unconventional myosins to filopodia functions.

## Material and methods

### Cells

U2-OS (human osteosarcoma) and MDA-MB-231 (triple-negative human breast adenocarcinoma) cells were grown in DMEM (Dulbecco’s Modified Eagle’s Medium with HEPES modification; Sigma, D1152) supplemented with 10 % fetal bovine serum (FCS) (Biowest, S1860). U2-OS cells were purchased from DSMZ (Leibniz Institute DSMZ-German Collection of Microorganisms and Cell Cultures, Braunschweig DE, ACC 785). CHO cells were cultured in alpha-MEM, supplemented with 5% FCS and L-glutamine. U2-OS, MDA-MB-231, and CHO cells were transfected using Lipofectamine 3000 and the P3000TM Enhancer Reagent (Thermo Fisher Scientific) according to the manufacturer’s instructions. The U2-OS MYO10-GFP lines were generated by transfecting U2-OS cells using Lipofectamine 3000 (ThermoFisher Scientific), selected using Geneticin (ThermoFisher Scientific; 400 μg.ml^-1^ final concentration) and sorted for green fluorescence using a fluorescence-assisted cell sorter (FACS). All cell lines tested negative for mycoplasma.

### Plasmids

EGFP-MYO10 was a gift from Emanuel Strehler (Addgene plasmid 47608) (Bennett et al., 2007). CRK-GFP was a gift from Ken Yamada (Addgene plasmid 50730). DIAPH3-GFP and VASP-GFP were gifts from Michael Davidson (Addgene plasmids 54158 and 54297, respectively). BCAR1-GFP was a gift from Daniel Rösel (Charles University in Prague, Czech Republic) (Braniš et al., 2017). FERMT2-GFP was a gift from Maddy Parsons (King’s College London, UK). The following constructs were described previously: GFP-ITGA2 and GFP-ITGA2^GAAKR^ (Pellinen et al., 2006), mScarlet-MYO10 (Jacquemet et al., 2019), GFP-TLN1 (Kopp et al., 2010), GFP-TLN1^FERM^ and His-TLN1^FERM^ (Goult et al., 2010).

The construct encoding the EGFP-tagged MYO10-FERM domain (EGFP-MYO10^FERM^) was designed using the boundaries from the MYO10-FERM crystal structure (Wei et al., 2011). The MYO10 coding region 1480-2053 was amplified by PCR (primers: 5’-ATT AGA GAA TTC AAC CCG GTG GTC CAG TGC-3’, 5’-ATT AGA GGT ACC TCA CCT GGA GCT GCC CTG-3’), and the resulting PCR products were ligated into pEGFP-C1 using the EcoRI and KpnI restriction sites. To generate the EGFP-MYO10-FERM^ITGBD^ mutant, a synthetic DNA sequence (gene block, IDT) encoding the MYO10 FERM domain (as indicated above) containing the appropriate mutations (S2001_F2002insA/T2009D) was inserted into pEGFP-C1 using the EcoRI/KpnI restriction sites. To generate the His-tagged MYO10^FERM^ plasmid, the MYO10-FERM domain (boundaries 1504-2058 in MYO10) was amplified by PCR (primers: 5’-ATT AGA GCG GCC GCA CCG ATC GAC ACC CCC AC, 5’-ATT AG AGA ATT CTC ACC TGG AGC TGC CCT G) and introduced in pET151 using the NotI and EcoRI restriction sites.

The MYO10 MyTH/FERM deletion construct (EGFP-MYO10^ΔF^) was generated by introducing a premature stop codon in full-length EGFP-MYO10 (boundaries 1-1512 in MYO10) using a gene block (IDT). The gene block was inserted in EGFP-MYO10 using the PvuI and XbaI restriction sites.

The mScarlet-I-MYO10^ΔF^ construct was created from EGFP-MYO10^ΔF^ by swapping the fluorescent tag. The mScarlet-I (Bindels et al., 2017) coding sequence, acquired as a gene block (IDT), was inserted in EGFP-MYO10^ΔF^ using the NheI and KpnI restriction sites.

The MYO10/TLN1 chimera construct (EGFP-MYO10^TF^) was generated by swapping the MYO10-FERM domain (boundaries 1504-2056 in MYO10) with the TLN1-FERM domain (boundaries 1-398 in TLN1) using a gene block (IDT). The gene block was inserted in EGFP-MYO10 using the PvuI and XbaI restriction sites.

The MYO10^ITGBD^ construct was generated by replacing the wild type MYO10-FERM domain (boundaries 1504-2056 in MYO10) with a MYO10 FERM domain containing the required mutations (S2001_F2002insA/T2009D) using a gene block (IDT). The gene block was inserted in EGFP-MYO10 using the PvuI and XbaI restriction sites.

The MYO10^ΔF2F3^ and MYO10^ΔF3^ constructs were generated by replacing the wild type MYO10-FERM domain (boundaries 1504-2056 in MYO10) with truncated MYO10 FERM domains where the F2-F3 or F3 FERM lobes are deleted using gene blocks (IDT). The gene blocks were inserted in EGFP-MYO10 using the PvuI and XbaI restriction sites. The final boundaries compared to full length MYO10 are 1-1794 for MYO10^ΔF2F3^ and 1-1951 for MYO10^ΔF3^. All the constructs generated in this study were validated by sequencing and are in the process of being deposited into Addgene.

### Antibodies and other reagents

Monoclonal antibodies recognizing the extended conformation of β1 integrin (high affinity for ligand, termed ‘active’) were mouse anti-human β1 integrin 12G10 (generated in house from a hybridoma), mouse anti-human β1 integrin HUTS21 (556048, BD Biosciences), and rat anti-human β1 integrin 9EG7 (BD Biosciences, 553715). Monoclonal antibodies recognizing the closed conformation of β1 integrin (unoccupied β1 integrin, termed ‘inactive’) were mouse anti-human β1 integrin 4B4 (Beckman Coulter, 6603113) and rat anti-human β1 integrin mAb13 (generated in house from a hybridoma). The monoclonal antibody recognizing all β1 integrin species was mouse anti-human β1 integrin P5D2 (Developmental studies hybridoma bank). Other mouse monoclonal antibodies used in this study were raised against hamster a5 integrin (antibody PB1, Developmental studies hybridoma bank), anti-human TLN1 (antibody 97H6, Novus Biologicals NBP2-50320), anti-human TLN2 (antibody 68E7, Novus Biologicals NBP2-50322), β-actin (antibody AC-15, Sigma, Cat. No. A1978) and PAX (antibody 349, BD Biosciences, 610051). The rabbit monoclonal antibody used was raised against AP2μ (Novus Biological, EP2695Y). Rabbit polyclonal antibodies used were raised against GFP (Abcam Ab290), MYO10 (Novus Biologicals, 22430002; 1:1000 for WB), and kindlin-1 (recognizes kindlin 1 and 2, Abcam, ab68041).

Small molecule inhibitors used were RO-3306 (CDK1 inhibitor, Sigma SML0569) and blebbistatin (Stemcell technologies 72402). The bovine plasma fibronectin was purchased from Merck (341631) and collagen I was purchased from Sigma (C8919-20ML).

### siRNA-mediated gene silencing

The expression of proteins of interest was suppressed using 83 nM siRNA and lipofectamine 3000 (Thermo Fisher Scientific) according to the manufacturer’s instructions. All siRNAs used were purchased from Qiagen. The siRNA used as control (siCTRL) was Allstars negative control siRNA (Qiagen, Cat No./ID: 1027280). siRNAs targeting ACTN1 were siACTN1 #5 (Hs_ACTN1_5, SI00299131) and siACTN1 #2 (Hs_ACTN1_2, SI00021917). siRNAs targeting TNS3 were siTNS3 #1 (Hs_TENS1_1, SI00134372) and siTNS3 #2 (Hs_TNS3_2, SI02778643). siRNAs targeting TNS1 were siTNS1 #3 (Hs_TNS_3, SI00134106) and siTNS1 #4 (Hs_TNS_4, SI00134113). siRNAs targeting FERMT1 were siFERMT1 #5 (Hs_C20orf42_5, SI04269181), siFERMT1 #7 (Hs_C20orf42_7, SI04307219) and siFERMT1 #8 (Hs_C20orf42_8, SI04352978). siRNAs targeting FERMT2 were siFERMT2 #1 (Hs_FERMT2_1, SI04952542) and siFERMT2 #3 (Hs_FERMT2_3, SI04952556). siRNAs targeting CIB1 were siCIB1 #5 (Hs_CIB1_5, SI02657102) and siCIB #7 (Hs_CIB1_7, SI03164476). siRNAs targeting SHARPIN were siSHARPIN #2 (Hs_SHARPIN_2, SI00140182) and siSHARPIN #5 (Hs_SHARPIN_5, SI03067344). siRNA targeting ITGB1BP1 were siITGB1BP1 #5 (Hs_ITGB1BP1_5, SI03129385) and siITGB1BP1 #8 (Hs_ITGB1BP1_8, SI04332832). siRNA targeting TLN1 were siTLN1 #2 (Hs_TLN1_2, SI00086968) and siTLN1 #3 (Hs_TLN1_3, SI00086975). siRNA targeting TLN2 was siTLN2 #3 (Hs_TLN2_3, SI00109277). siRNA targeting MYO10 were siMYO10 #5 (Hs_MYO10_5, SI04158245), siMYO10 #6 (Hs_MYO10_6, SI04252822) and siMYO10 #7 (Hs_MYO10_7, SI05085507). siMYO10 #7 targets the 3’ UTR of the MYO10 mRNA and therefore does not affect the expression of MYO10 constructs.

### SDS–PAGE and quantitative western blotting

Purified proteins or protein extracts were separated under denaturing conditions by SDS–PAGE and transferred to nitrocellulose membrane using Trans-Blot Turbo nitrocellulose transfer pack (Bio-Rad, 1704159). Membranes were blocked for 45 min at room temperature using 1x StartingBlock buffer (Thermo Fisher Scientific, 37578). After blocking, membranes were incubated overnight with the appropriate primary antibody (1:1000 in PBS), washed three times in TBST, and probed for 40 min using a fluorophore-conjugated secondary antibody diluted 1:5000 in the blocking buffer. Membranes were washed three times using TBST, over 15 min, and scanned using an Odyssey infrared imaging system (LI-COR Biosciences).

### siRNA screen

96-well glass-bottom plates (Cellvis, P96-1.5H-N) were first coated with a solution of poly-D-lysine (10 μg/ml in PBS, Sigma-Aldrich, A-003-M) at 4°C overnight. Plates were then washed with PBS and coated with a solution containing 10 μg/ml of bovine fibronectin (in PBS) also at 4°C overnight. Excess fibronectin was washed away with PBS.

U2-OS cells stably expressing MYO10-GFP were silenced for the gene of interest using a panel of siRNAs (Qiagen flexiplate, 1704159) using Lipofectamine 3000 (Thermo Fisher Scientific, L3000075). 48 h post silencing, cells were trypsinized and plated on both fibronectin-coated 96-well glass-bottom plates and 96-well plastic-bottom plates in full culture medium. Cells plated in the plastic-bottom plates were allowed to spread for two hours before being lysed using an RNA extraction buffer. RNAs were then purified and the silencing efficiency of each siRNA was validated by qPCR analysis.

Cells plated in the glass-bottom plates were allowed to spread for two hours and fixed with a warm solution of 4% paraformaldehyde (PFA; Thermo Scientific, 28906). After washing, the samples were incubated with a solution of 1 M glycine (30 min, in PBS) and then for one hour in a solution containing phalloidin–Atto647N (1/400 in PBS, Thermo Fisher Scientific, 65906) and DAPI (0.5 μg/ml in PBS, Thermo Fisher Scientific, D1306). The 96-well glass-bottom plates were then imaged using a spinning-disk confocal microscope equipped with a 40x objective. Images were analyzed using Fiji (Schindelin et al., 2012). Briefly, images were opened and, after background subtraction and normalization, MYO10 spots were automatically detected using Michael Schmid’s ‘Find maxima’ plugin. As inactive MYO10 is known to accumulate in rab7 vesicles (Plantard et al., 2010), to obtain an accurate number of filopodia-specific MYO10 spots, intracellular MYO10 spots were excluded from the analysis. Intracellular MYO10 spots were automatically filtered by masking the cells using the F-actin staining. The remaining spots per field of view were counted.

### RNA extraction, cDNA preparation, and Taq-Man qPCR

Total RNA extracted using the NucleoSpin RNA Kit (Macherey-Nagel, 740955.240C) was reverse transcribed into cDNA using the high-capacity cDNA reverse transcription kit (Applied Biosystems, Thermo Fisher Scientific, 43-688-14) according to the manufacturer’s instructions. The TaqMan primer sequences and associated universal probes were generated using ProbeFinder (version 2.53, Roche). The primers themselves were ordered from IDT, and the TaqMan fast advanced master mix (Thermo Fisher Scientific, 4444557) was used to perform the qPCR reactions according to the manufacturer’s instructions. Primers used in this study were against TNS1 (cca gac acc cac ctg act tag; ttg gtg cat tct cag tgg tg; probe 58), ACTN1 (gcc tca tca gct tgg gtt at; cat gat gcg ggc aaa ttc; probe 7), FERMT1 (aga cgt cac act gag agt atc tgg; tct gac cag tct tgg gat ata ttg; probe 25), TNS3 (agg ctg cct gac aca gga; ;agg ggc tgt tca gca gag; probe 57), TLN1 (ccc tta cct ggg gag aca at; gag ctc acg gct ttg gtg; probe 61), CIB1 (agt tcc agc acg tca tct cc; gct gct gtc aca gga caa tc; probe 17), ITGB1BP (ttg aag ggc cat tag acc tg; gaa caa aag gca act ttc cat c; probe 61), FERMT2 (taa aa cat ggc gtt tca gca; cat ctg caa act cta cgg tgac; probe 48), SHARPIN (ccc tgg ctg tga gat gtg ta; ggc cac tct ccc ctt gta ac; probe 83), FLNA (gtc acc ggt cgc tct cag; agg gga cgg ccc ttt aat; probe 32) and TLN2 (ggt cat ggt tgg gca gat; gca tgc ttg tgt tga tgg tc; probe 40). qPCR reactions were analyzed with the 7900HT fast RT-PCR System (Applied Biosystems), and the results were analyzed using the RQ Manager Software (Applied Biosystems). Relative expression was calculated by the 2^-ΔΔCT^ method. GAPDH mRNA levels were used to normalize data between experiments and conditions.

### Generation of filopodia maps

U2-OS cells transiently expressing the constructs of interests were plated on high tolerance glass-bottom dishes (MatTek Corporation, coverslip #1.7) pre-coated first with Poly-L-lysine (10 μg/ml, 1 h at 37°C) and then with bovine plasma fibronectin (10 μg/ml, 2 h at 37°C). After 2 h, samples were fixed and permeabilized simultaneously using a solution of 4% (wt/vol) PFA and 0.25% (vol/vol) Triton X-100 for 10 min. Cells were then washed with PBS, quenched using a solution of 1 M glycine for 30 min, and, when appropriate, incubated with the primary antibody for 1 h (1:100). After three washes, cells were incubated with a secondary antibody for 1 h (1:100). Samples were then washed three times and incubated with SiR-actin (100 nM in PBS; Cytoskeleton; catalog number: CY-SC001) at 4°C until imaging (minimum length of staining, overnight at 4°C; maximum length, one week). Just before imaging, samples were washed three times in PBS and mounted in vectashield (Vector Laboratories).

To map the localization of each protein within filopodia, images were first processed in Fiji (Schindelin et al., 2012) and data analyzed using R as previously described (Jacquemet et al., 2019). Briefly, in Fiji, the brightness and contrast of each image was automatically adjusted using, as an upper maximum, the brightest cellular structure labeled in the field of view. In Fiji, line intensity profiles (1-pixel width) were manually drawn from filopodium tip to base (defined by the intersection of the filopodium and the lamellipodium). To avoid any bias in the analysis, the intensity profile lines were drawn from a merged image. All visible filopodia in each image were analyzed and exported for further analysis (export was performed using the ‘‘Multi Plot’’ function). For each staining, line intensity profiles were then compiled and analyzed in R. To homogenize filopodia length; each line intensity profile was binned into 40 bins (using the median value of pixels in each bin and the R function ‘‘tapply’’). Using the line intensity profiles, the percentage of filopodia positive for active β1 at their tip was quantified. A positive identification was defined as requiring at least an average value of 5000 (values between 0-65535) within the bins defining the filopodium tip (identified using MYO10 staining). The map of each protein of interest was created by averaging hundreds of binned intensity profiles. The length of each filopodia analyzed was directly extracted from the line intensity profiles.

The preferential recruitment of active and inactive β1 integrin to filopodia tips or shafts was assessed by calculating an enrichment ratio where the averaged intensity of the β1 integrin species at the filopodium tip (bin 1-6) was divided by the averaged intensity at the filopodium shaft (bin 7-40). This enrichment ratio was calculated for each filopodium analyzed and the results displayed as Tukey box plots.

### Quantification of filopodia numbers and dynamics

For the filopodia formation assays, cells were plated on fibronectin-coated glass-bottom dishes (MatTek Corporation) for 2 h. Samples were fixed for 10 min using a solution of 4% PFA, then permeabilized using a solution of 0.25% (vol/vol) Triton X-100 for 3 min. Cells were then washed with PBS and quenched using a solution of 1 M glycine for 30 min. Samples were then washed three times in PBS and stored in PBS containing SiR-actin (100 nM; Cytoskeleton; catalog number: CY-SC001) at 4°C until imaging. Just before imaging, samples were washed three times in PBS. Images were acquired using a spinning-disk confocal microscope (100x objective). The number of filopodia per cell was manually scored using Fiji (Schindelin et al., 2012).

To study filopodia stability, U2-OS cells expressing MYO10-GFP were plated for at least 2 h on fibronectin before the start of live imaging (pictures taken every 5 s at 37°C, on an Airyscan microscope, using a 40x objective). All live-cell imaging experiments were performed in normal growth media, supplemented with 50 mM HEPES, at 37°C and in the presence of 5% CO_2_. Filopodia lifetimes were then measured by identifying and tracking all MYO10 spots using the Fiji plugin TrackMate (Tinevez et al., 2017). In TrackMate, the LoG detector (estimated bob diameter = 0.8 mm; threshold = 20; subpixel localization enabled) and the simple LAP tracker (linking max distance = 1 mm; gap-closing max distance = 1 mm; gap-closing max frame gap = 0) were used.

### Light microscopy setup

The spinning-disk confocal microscope (spinning-disk confocal) used was a Marianas spinning-disk imaging system with a Yokogawa CSU-W1 scanning unit on an inverted Zeiss Axio Observer Z1 microscope controlled by SlideBook 6 (Intelligent Imaging Innovations, Inc.). Images were acquired using either an Orca Flash 4 sCMOS camera (chip size 2,048 × 2,048; Hamamatsu Photonics) or an Evolve 512 EMCCD camera (chip size 512 × 512; Photometrics). Objectives used were a 40x water (NA 1.1, LD C-Apochromat, Zeiss), a 63× oil (NA 1.4, Plan-Apochromat, M27 with DIC III Prism, Zeiss) and a 100x oil (NA 1.4 oil, Plan-Apochromat, M27) objective.

The structured illumination microscope (SIM) used was DeltaVision OMX v4 (GE Healthcare Life Sciences) fitted with a 60x Plan-Apochromat objective lens, 1.42 NA (immersion oil RI of 1.516) used in SIM illumination mode (five phases x three rotations). Emitted light was collected on a front-illuminated pco.edge sCMOS (pixel size 6.5 mm, readout speed 95 MHz; PCO AG) controlled by SoftWorx.

The confocal microscope used was a laser scanning confocal microscope LSM880 (Zeiss) equipped with an Airyscan detector (Carl Zeiss) and a 40x oil (NA 1.4) objective. The microscope was controlled using Zen Black (2.3), and the Airyscan was used in standard super-resolution mode.

### Integrin activity assays

CHO cells detached using Hyclone HyQTase (Thermo Fisher Scientific, SV300.30.01), washed with Tyrode’s Buffer (10 mM HEPES-NaOH, pH 7.5, 137 mM NaCl, 2.68 mM KCl, 0.42 mM NaH_2_PO_4_, 1.7 mM MgCl_2_, 11.9 mM NaHCO_3_, 5 mM glucose, and 0.1% BSA) and pretreated for 10 min with or without 5 mM EDTA in serum-free alpha-MEM media. Cells were then incubated for 40 min with Alexa Fluor 647 labeled fibronectin fragment (FN 7-10). After washing away the unbound fibronectin using Tyrode’s buffer, cells were fixed with 4 % PFA (in PBS) for 10 min at room temperature. Part of the HyQTase treated cells were also fixed with 4 % PFA (in PBS) and stained with an anti-hamster anti-α5 integrin antibody to detect total ITGA5 levels in cells (2 h at 4C, 1:10 in PBS, antibody PB1, Developmental studies hybridoma bank) and with an Alexa Fluor 647-conjugated secondary antibody (45 min at RT, 1:200 in PBS, Thermo Fisher Scientific, A-21235). Fluorescence intensity was recorded using FACS (BD LSRFortessa™). Data were gated and analyzed using the Flowing Software (http://flowingsoftware.btk.fi/). The integrin activity index (IA) was calculated for each condition as a ratio AI = (F−F_EDTA_)/(F_PB1_), where F = FN7-10 signal, F_EDTA_ = FN7-10 signal in EDTA treated cells and F_PB1_ = a5 integrin signal.

MDA-MB-231 and U2-OS cells detached using Hyclone HyQTase (Thermo Fisher Scientific, SV300.30.01) were fixed with 4 % PFA (in PBS) for 10 min and stained for active (antibody 9EG7) and total β1 integrin (antibody P5D2) overnight at 4°C. Cells were then stained with the appropriate Alexa Fluor 647-conjugated secondary antibody (45 min at RT, 1:200, Thermo Fisher Scientific) and the fluorescence was recorded using FACS. Data were gated and analyzed using the Flowing Software (http://flowingsoftware.btk.fi/) and the integrin activity (IA) was calculated as indicated below where F_9EG7_ and F_P5D2_ are the signals intensities of the 9EG7 and P5D2 stainings, respectively. F_2nd Ab_ corresponds to the signal intensity recorded when the cells are stained with only the secondary antibody.

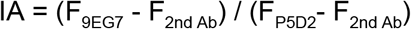

### Cell spreading assay

The xCELLigence RTCA instrument (Roche) was used to measure cell adhesion on fibronectin in real-time (Hamidi et al., 2017). The RTCA instrument uses gold-bottom electrode plates to measure the impedance between two electrodes. This is expressed as an arbitrary cell index value. The xCELLigence 96-well plates (Acea Biosciences, E-Plate VIEW 96 PET, 00300600900) were coated with a solution of 20 μg/ml of poly-D-lysine (in PBS) for 1 h at 37°C, washed with PBS, and coated with a solution of 10 μg/ml fibronectin (in PBS) for 1 h at 37°C. Plates were then blocked using a solution of 1% BSA (in PBS) for 1 h in RT. After 2 PBS washes, 15000 cells were seeded into each well in a serum-free culture medium. The cell index was recorded over time.

### Recombinant protein expression and purification

The *E. coli* BL-21(DE3) strain was transformed with IPTG inducible, His-tagged expression constructs, and the transformed bacteria were grown at 37°C in LB media supplemented with ampicillin (1 mg/ml) until OD600 was 0.6-0.8. Protein expression was then induced using IPTG (0.5 mM), and the temperature was lowered to 25°C. Cells were harvested after 5 h by centrifugation (20 min at 6000 g). Bacteria were then resuspended in a resuspension buffer (1x TBS, cOmplete™ protease inhibitor tablet (Roche, cat. no. 5056489001), 1x AEBSF inhibitor, 1x PMSF, RNase 0.05 mg/ml, DNase 0.05 mg/ml). To lyse the bacteria, a small spoonful of lysozyme and 1x BugBuster (Merck Millipore, cat. no. 70584-4) were added, and the suspension was agitated for 30 min at 4°C. Cell debris was pelleted using a JA25.5 rotor at 20000 rpm for 1 h. His-tagged proteins were batch purified from the supernatant using a Protino Ni-TED 2000 column (Macherey Nagel, cat. no. 745120.25) according to the manufacturer’s instructions. Proteins were eluted using the elution buffer provided with the kit supplemented with 1 mM AEBSF. For each purified protein, several 1 ml fractions were collected, ran on a 4-20 % protein gel (Bio-Rad Mini-PROTEAN TGX, #4561093), stained with InstantBlue® (Expedeon, ISB1L), and the fractions abundant in tagged protein were combined. Imidazole was removed in a buffer exchange overnight at 4°C and 1 mM AEBSF was added to the imidazole-free protein. Proteins were stored at 4°C for up to one week.

### Whole-mount immuno-SEM

U2-OS cells expressing MYO10-GFP were plated for 2 h on fibronectin-coated coverslips and fixed with a solution of 4 % PFA (in 0.1 M HEPES, pH 7.3) for 30 min. After washing and quenching with 50 mM NH_4_Cl (in 0.1 M HEPES), non-specific binding was blocked with a buffer containing 2 % BSA (in 0.1 M HEPES). Samples were then labeled using the appropriate primary antibody (1:10 in 0.1 M HEPES) for 30 min, washed, and labeled with a gold conjugated secondary antibody (1:50 in 0.1 M HEPES, 30 nm gold particles, BBI solutions, EM.GAF30) for 30 min. After immunolabeling, the samples were washed, and post-fixed with a solution of 2.5 % glutaraldehyde and 1 % buffered osmium tetroxide prior to dehydration and drying using hexamethyldisilazane. The dried samples were mounted on SEM stubs and sputter-coated with carbon. The micrographs were acquired with FEI Quanta FEG 250 microscope with SE and vC detectors (FEI Comp.) using an acceleration voltage of 5.00 kV and a working distance ranging from 7.7 to 10.9 mm.

### Integrin tail pull-downs

For each pulldown, 20 μl of streptavidin Dynabeads (MyOne Streptavidin C1, Invitrogen, 65001) were incubated, for 30 min, on ice, with the appropriate biotinylated integrin tail peptides (50 ug per sample) (LifeTein). U2-OS cells were washed twice with cold PBS and lysed on ice with a buffer containing 40 mM HEPES, 75 mM NaCl, 2 mM EDTA, 1 % NP-40, a cOmplete™ protease inhibitor tablet (Roche, 5056489001) and a phosphatase-inhibitor tablet (Roche, 04906837001). Samples were cleared by centrifugation (13,000 g, 10 min) and incubated with the streptavidin Dynabeads for 2 h at 4°C. Beads were washed three times with a washing buffer (50 mM Tris-HCl pH 7.5, 150 mM NaCl, 1 % (v/v) NP-40), and proteins bound to the beads were eluted using SDS sample buffer and heated for 5-10 min at 90°C. Results were analyzed using western blot. Integrin peptides used were wild-type β1-integrin tail (KLLMIIHDRREFAKFEKEKMNAKWDTGENPIYKSAVTTVVNPKYEGK), the β1-integrin tail where the NPXY motif is deleted (KLLMIIHDRREFAKFEKEKMNAKWDTGEN), the conserved region of the α2-integrin tail (WKLGFFKRKYEKM), the conserved region of α2-integrin tail peptide where the GFFKR motif is mutated (GAAKR mutant, WKLGAAKRKYEKM) and the wild-type α5-integrin tail (KLGFFKRSLPYGTAMEKAQLKPPATSDA).

### Microscale thermophoresis

Recombinant His-tagged proteins were labeled using the Monolith His-Tag Labeling Kit RED-tris-NTA (NanoTemper, MO-L008). In all experiments, the labeled His-tagged recombinant proteins were used at a concentration of 20 nM while the integrin tail peptides were used at increasing concentration. Kd values were calculated using the equation provided below (Eq.1), where Kd is the dissociation constant, [A] the concentration of the free fluorescent molecule, [L] the concentration of the free ligand, [AL] the concentration of the AL-complex. [A0] is the known concentration of the fluorescent molecule and [L0] is the known concentration of added ligand. This leads to a quadratic fitting function for [AL]:

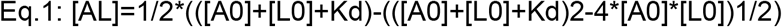

Alternatively, binding was also expressed as a change in MST signal (normalized fluorescence ΔFnorm). This is defined as a ratio:

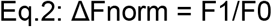

Where F0 is the fluorescence prior and F1 after IR laser activation.

All binding data were analyzed using MO.Control and MO.Affinity software (NanoTemper).

### Quantification and statistical analysis

Randomization tests were performed using the online tool PlotsOfDifferences (https://huygens.science.uva.nl/PlotsOfDifferences/) (Goedhart, 2019). Dot plots were generated using PlotsOfData (Postma and Goedhart, 2019). SuperPlots were generated using SuperPlotsofData (Lord et al., 2020; Goedhart, 2020). Bar plots with visualized data points, time-series data, and density plots were generated using R (https://www.r-project.org/), Rstudio (Integrated Development for R. RStudio, Inc., Boston, MA. http://www.rstudio.com/) and ggplot2 (Wickham, 2016). Other statistical analyses were performed using Google sheets except for the one-sample t-test which was performed using an online calculator (https://www.socscistatistics.com/tests/tsinglesample/default.aspx).

## Data availability

The authors declare that the data supporting the findings of this study are available within the article and from the authors on request.

## Acknowledgments

We thank K. Baker for technical assistance and the generation of the MYO10 His-tagged construct. We thank J. Siivonen, P. Laasola, J. Conway, and C. Guzmán for technical assistance, M. Saari for help with the microscopes, and H. Hamidi for editing and critical assessment of the manuscript. We thank Maria Taskinen, S. Salomaa, G. Follain, and H. Al Akhrass for providing comments on the manuscript. We thank E. Strehler, K. Yamada, M. Davidson, D. Rösel, and M. Parsons for providing reagents. The Cell Imaging and Cytometry Core facility (Turku Bioscience, University of Turku, Åbo Akademi University and Biocenter Finland) and Electron microscopy unit (Institute of Biotechnology, University of Helsinki) are acknowledged for services, instrumentation, and expertise. Both imaging units are supported by Biocenter Finland.

## Author contributions

Conceptualization, G.J. and J.I.; Methodology, M.M., H.V., E.J., B.T.G., J.I. and G.J.; Formal Analysis, M.M., M.L.B.G. and G.J.; Investigation, M.M., M.L.B.G., H.V., E.J., B.T.G., J.I. and G.J.; Writing – Original Draft, M.M., J.I. and G.J.; Writing – Review and Editing, M.M., M.L.B.G., H.V., E.J., B.T.G., J.I. and G.J.; Visualization, M.M., J.I. and G.J.; Supervision, G.J. and J.I.; Funding Acquisition, G.J. and J.I.

---

**Figure S1:**
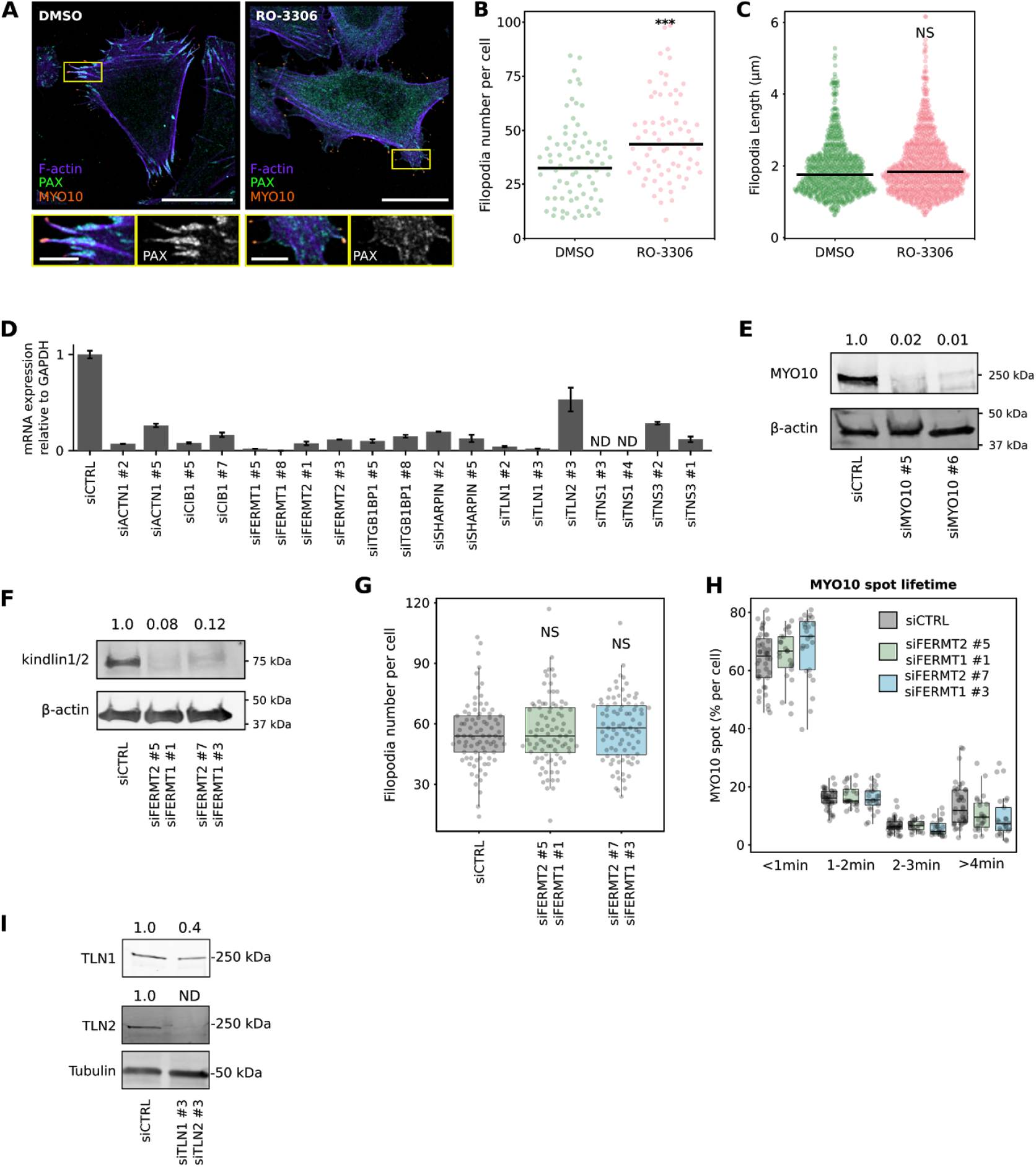
Modulation of filopodia properties by focal adhesions and known integrin activity regulators. **(A-C**) U2-OS cells expressing EGFP-MYO10 were plated on fibronectin for 1 h and treated for another hour with 10 μM RO-3306 (CDK1 inhibitor) or DMSO. Cells were stained for paxillin (PAX) and F-actin and imaged using an Airyscan confocal microscope or a spinning-disk confocal microscope. (**A**) Representative Airyscan images are displayed. The yellow rectangles highlight ROIs, which are magnified; scale bars: (main) 25 μm; (inset) 5 μm. (**B**) The number of MYO10-positive filopodia per cell was then quantified from the spinning-disk images (n > 72 cells, two biological repeats; *** p-value = 0.003). **C**) Quantification of filopodia length, from SIM images, in U2-OS cells transiently expressing EGFP-MYO10 and treated for 1 h with 10 μM RO-3306 (CDK1 inhibitor) or DMSO (DMSO, n = 734 filopodia; RO-3306, n = 824 filopodia; three biological repeats; *** p-value = <0.001). **(D**) The efficiency of siRNA-mediated silencing of each target of the siRNA screen performed in Fig 2A (except MYO10) was quantified by qPCR and normalized to GAPDH expression. The results were further normalized against expression detected in siCTRL cells. **(E**) Efficiency of siRNA-mediated silencing of MYO10 (oligos #5 and #6) in U2-OS cells validated by western blot. (**F**) Efficiency of dual siRNA-mediated silencing of FERMT1 and FERMT2 in U2-OS cells validated by western blot. (**G**) FERMT1- and FERMT2-silenced U2-OS cells transiently expressing EGFP-MYO10 were plated on fibronectin for 2 h, fixed, and the number of MYO10-positive filopodia per cell was quantified (n > 70 cells, three biological repeats). (**H**) FERMT1- and FERMT2-silenced U2-OS cells transiently expressing EGFP-MYO10 were plated on fibronectin and imaged live using an Airyscan confocal microscope (1 picture every 5 s over 20 min). For each condition, MYO10-positive particles were automatically tracked, and MYO10 spot lifetime (calculated as a percentage of the total number of filopodia generated per cell) was plotted and displayed as boxplots (see Methods for details; three biological repeats, more than 21 cells per condition). **I**) Efficiency of dual siRNA-mediated silencing of TLN1 and 2 (oligos #3 and #3) in U2-OS cells (one round of silencing) validated by western blot. For all panels, p-values were determined using a randomization test. NS indicates no statistical difference between the mean values of the highlighted condition and the control.

**Figure S2.**
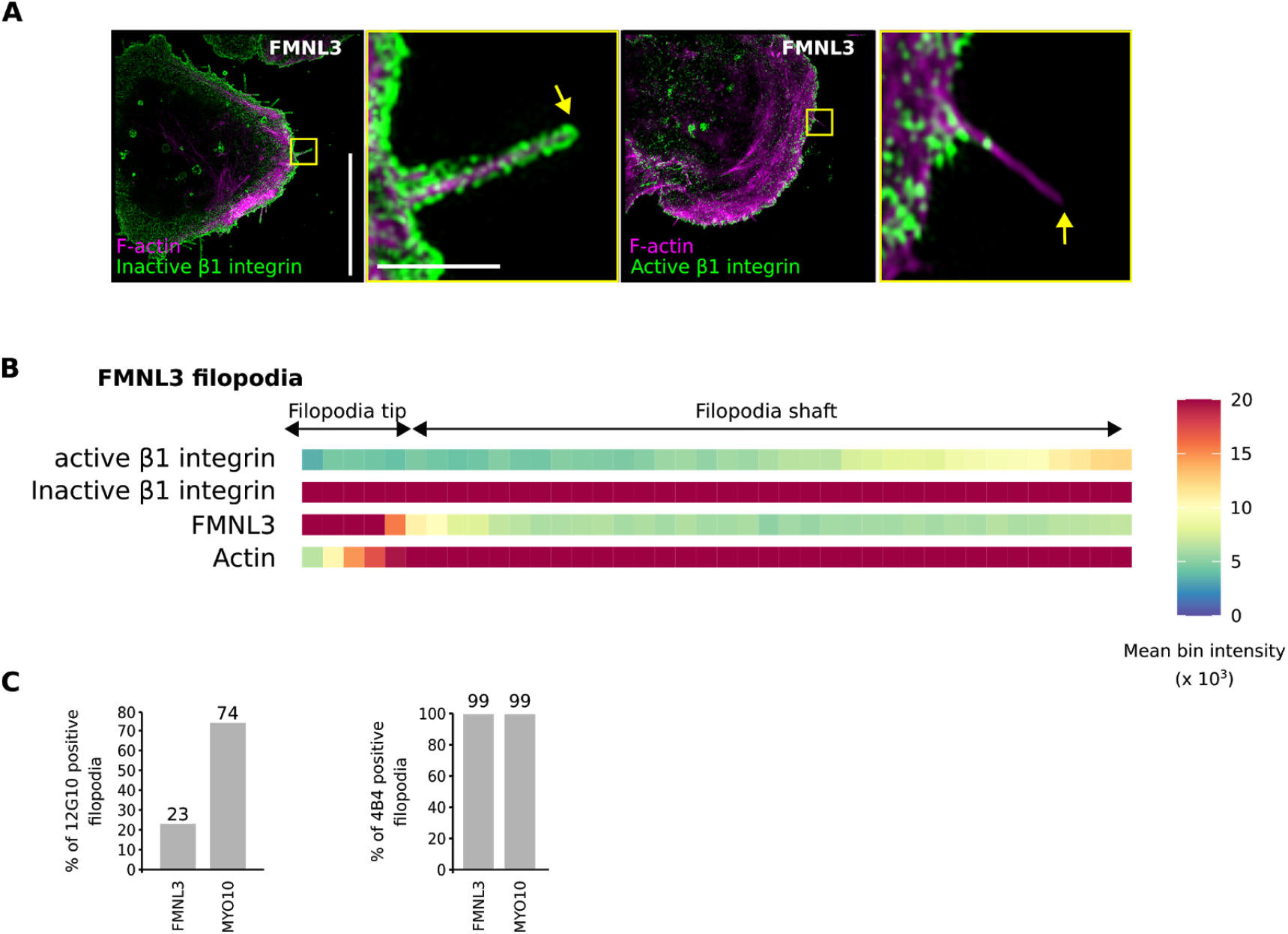
FMNL3-induced filopodia lack active integrin at their tips. **(A-C**) U2-OS cells expressing EGFP-FMNL3 were plated on fibronectin for 2 h, stained for active (antibody 12G10) or inactive (antibody mAb13) β1 integrin and F-actin, and imaged using SIM. **A**) Representative MIPs are displayed. The yellow squares highlight ROIs, which are magnified; yellow arrows highlight filopodia tips; scale bars: (main) 20 μm; (inset) 2 μm. **(B**) Heatmap highlighting the sub-filopodial localisation of the proteins stained in A based on their intensity profiles (FMNL3, n = 373 filopodia; F-actin, n = 373 filopodia; active β1 integrin, n = 228 filopodia; inactive β1 integrin, n = 143 filopodia). **(C**) Bar chart highlighting the percentage of FMNL3 and MYO10-induced filopodia with detectable levels of active (12G10) and inactive (4B4) β1 integrin (12G10: FMNL3, n = 228 filopodia; MYO10, n = 329 filopodia; 4B4: FMNL3, n = 143 filopodia; MYO10, n = 413 filopodia).

**Figure S3.**
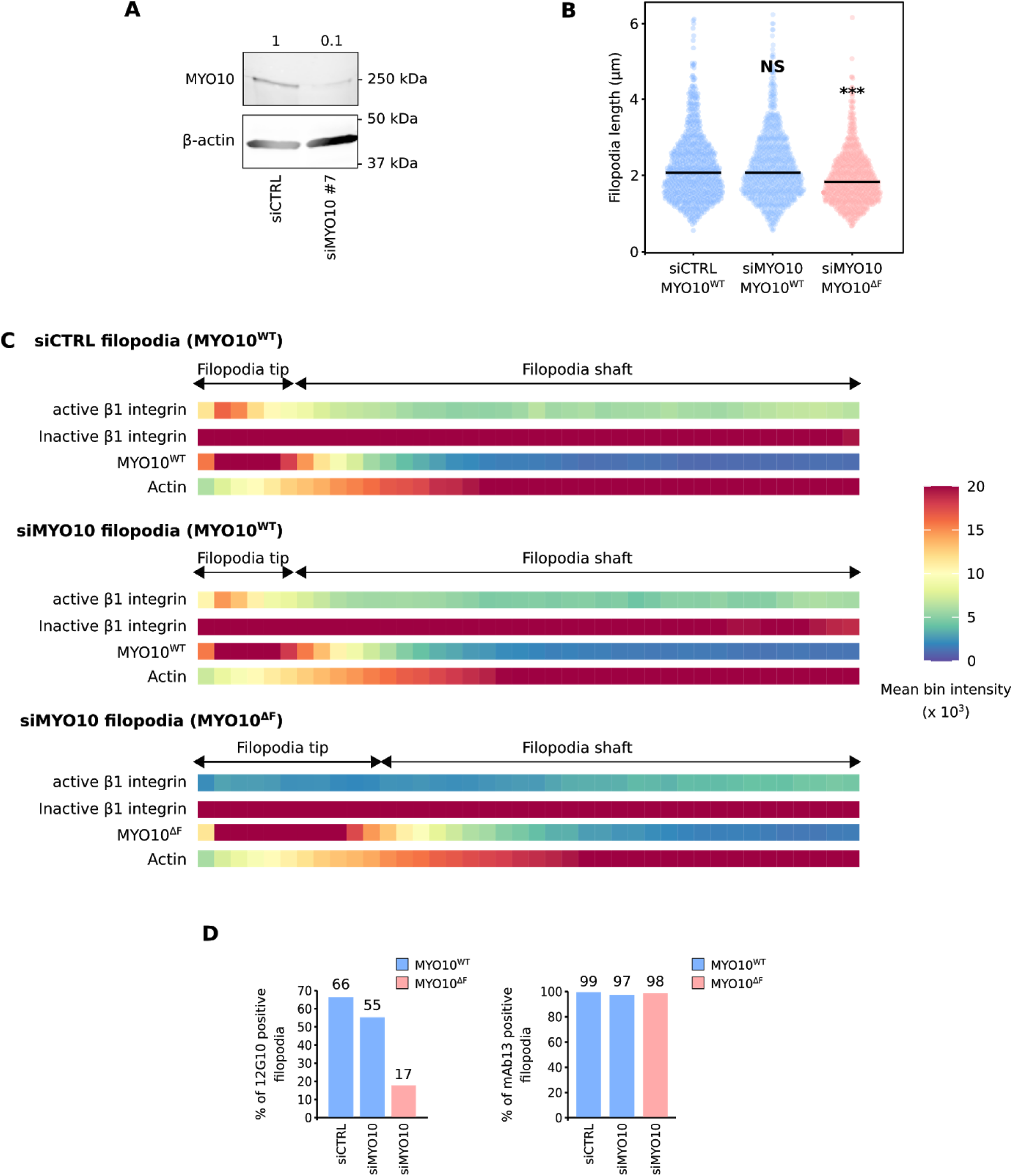
MYO10-FERM is required for integrin activation but not localization in filopodia. **(A**) The efficiency of siRNA-mediated silencing of MYO10 in U2-OS cells was validated by western blot. The siMYO10 #7 oligo targets the 3’ UTR of the MYO10 mRNA. **(B-D**) MYO10-silenced U2-OS cells transiently expressing EGP-MYO10^WT^ or MYO10^ΔF^ were plated on fibronectin, stained for active (antibody 12G10) or inactive (antibody mAb13) β1 integrin and F-actin, and imaged using SIM. **(B)** Quantification of filopodia length, from the SIM images, are displayed as dotplots where the median is highlighted (siCTRL EGP-MYO10^WT^, n = 799 filopodia; siMYO10 #7 EGP-MYO10^WT^, n = 897 filopodia; siMYO10 #7 EGP-MYO10^ΔF^, n = 731 filopodia; three biological repeats; *** p value = <0.001). **(C**) Heatmap highlighting the sub-filopodial localisation of the indicated proteins based on their intensity profiles (siCTRL EGP-MYO10^WT^ filopodia: MYO10, n = 799 filopodia; F-actin, n = 799 filopodia; active β1 integrin, n = 799 filopodia; inactive β1 integrin, n = 878 filopodia. siMYO10 #7 EGP-MYO10^WT^ filopodia: MYO10, n = 897 filopodia; F-actin, n = 897 filopodia; active β1 integrin, n = 897 filopodia; inactive β1 integrin, n = 960 filopodia. siMYO10 #7 EGP-MYO10^ΔF^ filopodia: MYO10, n = 731 filopodia; F-actin, n = 731 filopodia; active β1 integrin, n = 731 filopodia; inactive β1 integrin, n = 778 filopodia. Three biological repeats). **(D**) Bar chart highlighting the percentage of filopodia with detectable levels of active and inactive β1 integrin in the indicated conditions (active β1 integrin: siCTRL EGP-MYO10^WT^ filopodia, n = 799 filopodia; siMYO10 #7 EGP-MYO10^WT^, n = 897 filopodia; siMYO10 #7 EGP-MYO10^ΔF^ filopodia, n = 731 filopodia. Inactive β1 integrin: siCTRL EGP-MYO10^WT^ filopodia, n = 878 filopodia; siMYO10 #7 EGP-MYO10^WT^, n = 960 filopodia; siMYO10 #7 EGP-MYO10^ΔF^ filopodia, n = 778 filopodia. Three biological repeats). For all panels, p-values were determined using a randomization test. NS indicates no statistical difference between the mean values of the highlighted condition and the control.

**Figure S4.**
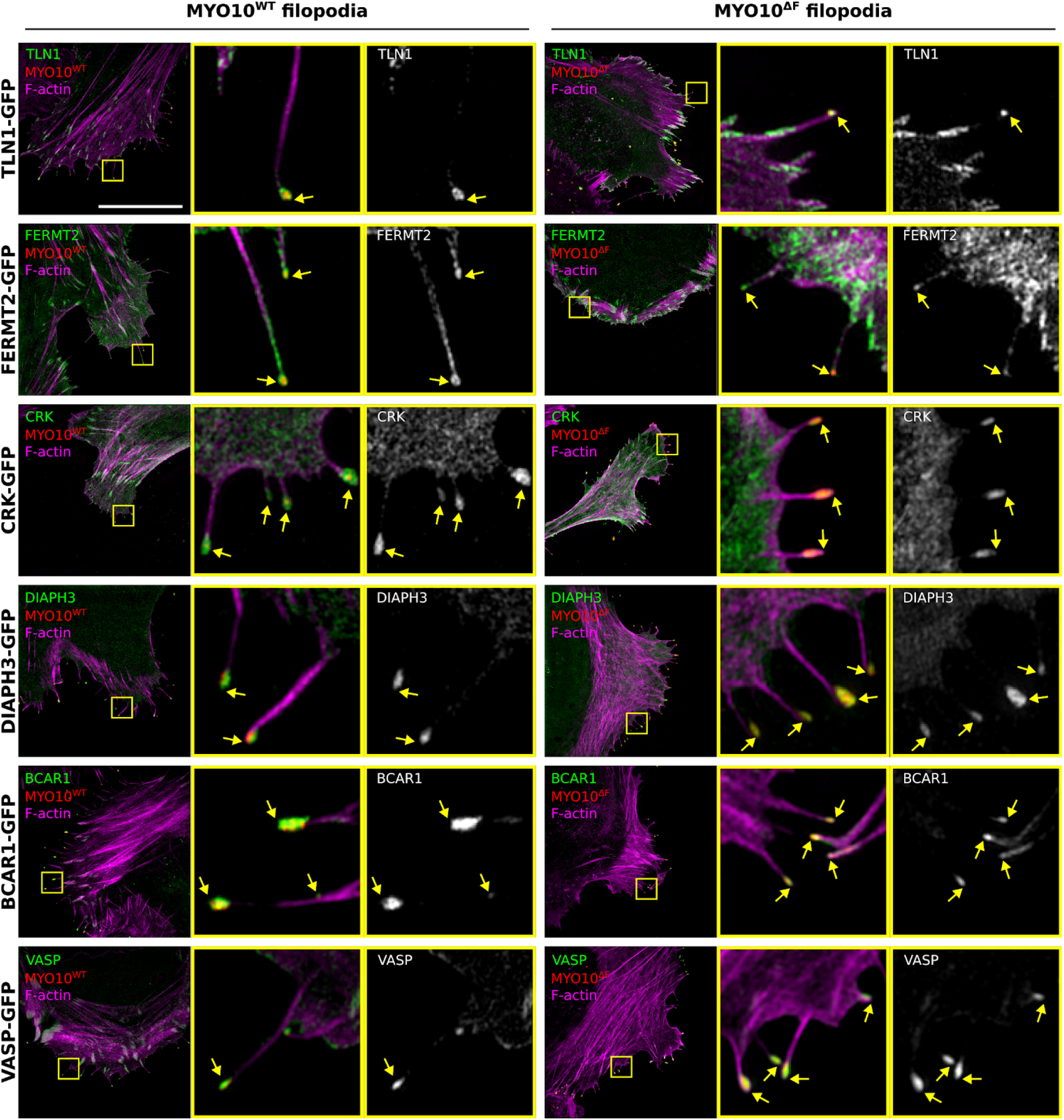
MYO10-FERM deletion has minimal impact on known filopodia tip protein localization. U2-OS cells expressing MYO10^WT^-mScarlet or MYO10^ΔF^-mScarlet-I together with TLN1-GFP, FERMT2-GFP, CRK-GFP, DIAPH3-GFP, BCAR1-GFP or VASP-GFP were plated on fibronectin for 2 h, fixed, stained for f-actin and imaged using SIM. Representative MIPs are displayed. The yellow squares highlight ROIs, which are magnified; yellow arrows highlight filopodia tips; scale bars: (main) 20 μm; (inset) 2 μm.

**Figure S5.**
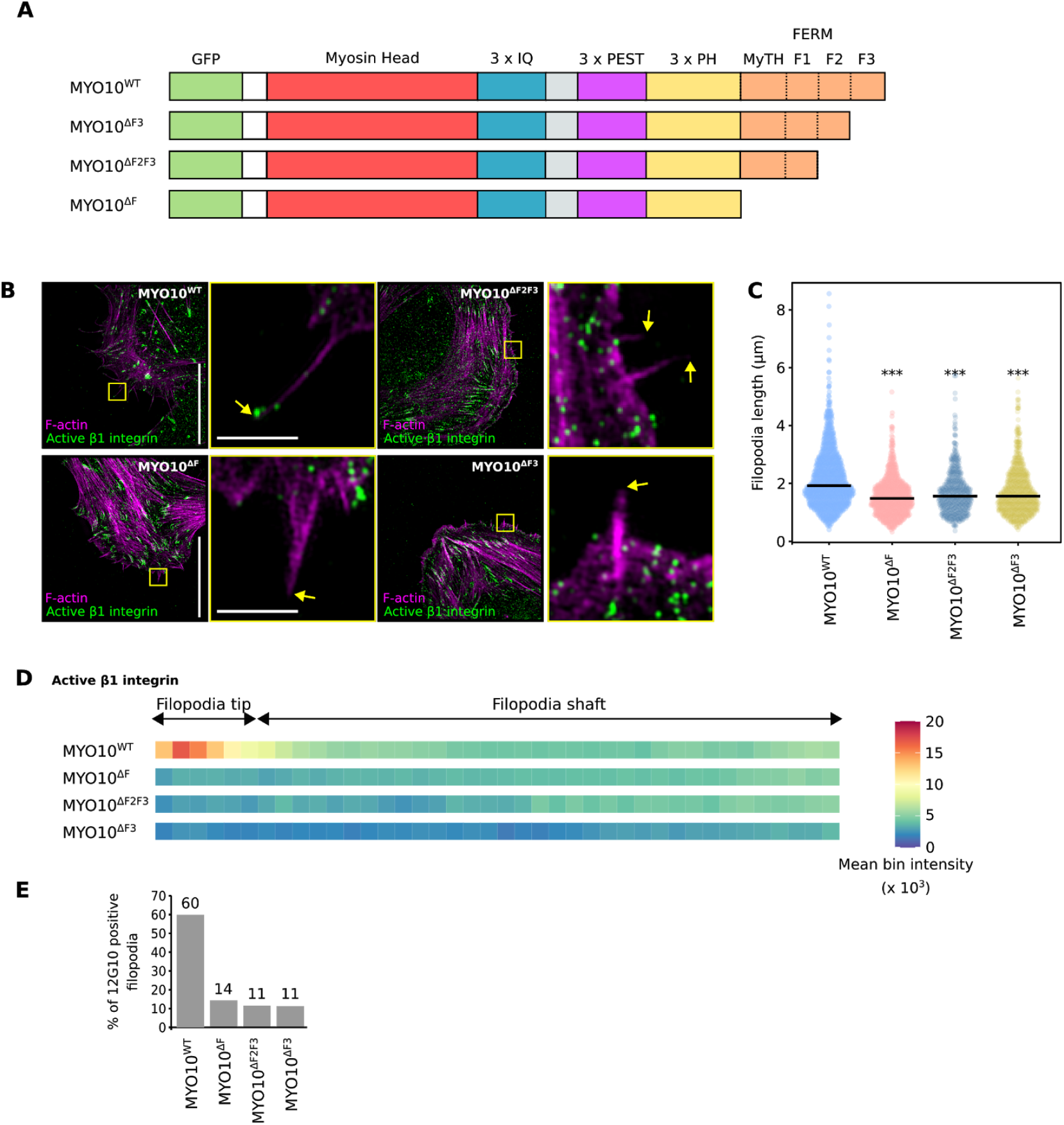
the F3 subdomain of MYO10-FERM is required to activate integrins at filopodia tips. **(A**) Cartoon illustrating the EGFP-MYO10^WT^, EGFP-MYO10^ΔF^, EGFP-MYO10^ΔF2F3^, and EGFP-MYO10^ΔF3^ constructs. (**B-E**) U2-OS cells transiently expressing EGFP-MYO10^WT^, EGFP-MYO10^ΔF^, EGFP-MYO10^ΔF2F3^ or EGFP-MYO10^ΔF3^ were plated on fibronectin, stained for active β1 integrin (antibody 12G10) and F-actin, and imaged using SIM. (**B**) Representative MIPs are displayed. The yellow squares highlight ROIs, which are magnified; yellow arrows highlight filopodia tips; scale bars: (main) 20 μm; (inset) 2 μm. (**C)** Quantification of MYO10^WT^, MYO10^ΔF^, MYO10^ΔF2F3^ and MYO10^ΔF3^ filopodia length, from the SIM images, are displayed as dot plots where the median is highlighted (*** p-value = <0.001). (**D**) Heatmap highlighting the sub-filopodial localization of active β1 integrin (antibody 12G10) in MYO10^WT^, MYO10^ΔF^, MYO10^ΔF2F3^ and MYO10^ΔF3^ filopodia. (**E**) Bar chart highlighting the percentage of MYO10^WT^, MYO10^ΔF^, MYO10^ΔF2F3^ and MYO10^ΔF3^ filopodia with detectable levels of active β1 integrin. (**B-E**) EGP-MYO10^WT^, n = 1073 filopodia; EGFP-MYO10^ΔF^, n = 776 filopodia; MYO10^ΔF2F3^, n = 497 filopodia; MYO10^ΔF3^, n = 723 filopodia; Three biological repeats. For all panels, p-values were determined using a randomization test.

**Figure S6.**
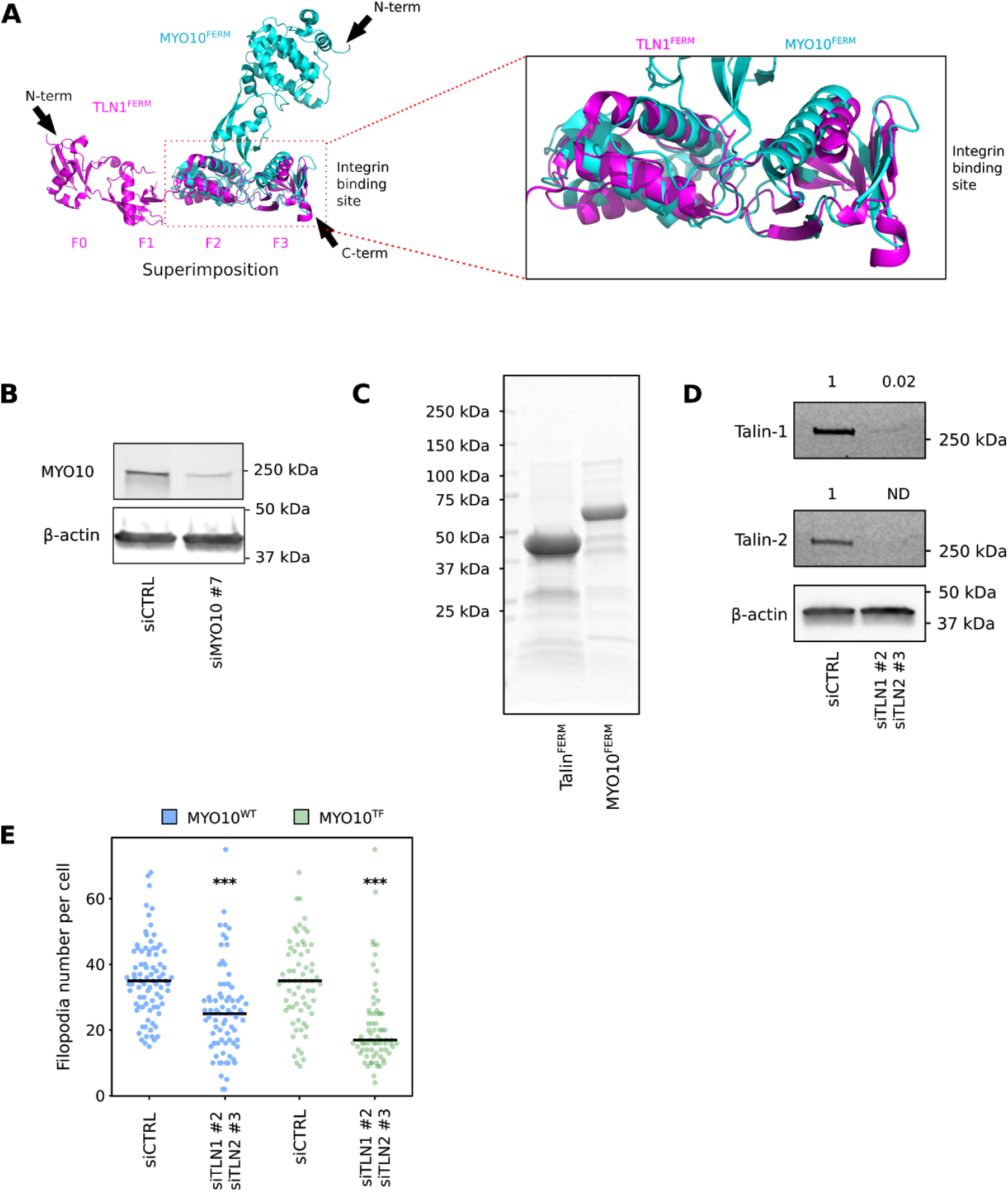
MYO10-FERM and talin-FERM structure and function in filopodia. **(A**) Visualisation of the structure of MYO10-FERM (PDB: 3PZD; (Wei et al., 2011)) and TLN1-FERM (PDB: 3IVF, (Elliott et al., 2010)) domains using PyMOL. The black arrows indicate the protein orientation from N to C terminal. The two FERM domains were superimposed to highlight their structural homology and differences. The integrin-binding region on the talin-FERM domain is highlighted and magnified. **(B)** The efficiency of siRNA-mediated silencing of MYO10 in MDA-231 cells was validated by western blot. The siMYO10 #7 oligo targets the 3’ UTR of the MYO10 mRNA. **(C**) Recombinant his-tagged TLN1 and MYO10-FERM domains were produced in bacteria and subsequently purified using a gravity Ni^2+^-column. A representative gel stained with Instant blue is displayed. **(D**) Efficiency of dual siRNA-mediated silencing of TLN1 and TLN2 in U2-OS cells following two rounds of silencing. A representative western blot is displayed. **(E**) TLN1 and TLN2-silenced U2-OS cells transiently expressing EGFP-MYO10^WT^ or EGFP-MYO10^TF^ were plated on fibronectin for 2 h, fixed, stained, imaged using a spinning-disk confocal microscope, and the number of MYO10-positive filopodia per cell was quantified (siCTRL/MYO10^WT^, n = 83 cells; siCTRL/MYO10^TF^, n = 66 cells; siTLN1 and siTLN2/MYO10^WT^, n = 75 cells; siTLN/MYO10^TF^, n = 71 cells; three biological repeats, *** p-value < 0.001). For all panels, p-values were determined using a randomization test.

